# Ribosomal L12 stalks recruit elongation factors to speed protein synthesis in *Escherichia coli*

**DOI:** 10.1101/2023.04.14.536948

**Authors:** Jennifer L. Hofmann, Theodore S. Yang, Alp M. Sunol, Roseanna N. Zia

## Abstract

Translating ribosomes must wait after each elongation step for a new ternary complex (EF-Tu·aa-tRNA·GTP) to arrive, facilitating rapid codon recognition testing. We recently showed that this wait-time rate-limits elongation in *Escherichia coli* due to competitive combinatoric searching through crowded cytoplasm by thousands of *E. coli*’s 42 unique ternary complexes. Here, we investigate whether ribosomal L12 subunits pool translation molecules to reduce this wait time. We mimic transport and reactions underlying elongation in a physiologically accurate, physically-resolved model of crowded cytoplasm. We find that L12 pre-loading as much as doubles translation rate by reducing diffusive search time. But more L12 is not always better: faster-growing bacteria tend to have fewer L12. We resolve this apparent contradiction by demonstrating tradeoffs between binding and novel sampling as a function of copy number in *E. coli*. Variable L12 copy numbers may thus have evolved for fast or slow bacterial growth as complementary survival strategies.

## INTRODUCTION

All cells must synthesize proteins to grow and divide. In fact, the translation process that produces proteins is the rate-limiting step of bacterial growth [1, 2]. Translation is carried out by the ribosome, which hosts the reaction of multiple translation molecules and initiation, elongation, and termination of protein synthesis. While reactions *inside* the ribosome are governed by well-established kinetics [3–7], the ‘wait time’ for codon recognition testing rate-limits elongation and can vary with growth rate and corresponding changes in cytoplasmic conditions [8]. This wait time is set by physical transport outside the ribosome, specifically a combinatoric competition between EF-Tu·aa-tRNA·GTP ternary complexes diffusing through crowded cytoplasm to the ribosome. The consequence of these physical effects — that elongation is rate-limited by biomolecule size, combinatorics, and crowding — cannot be adequately captured with adjustments to kinetic parameters or scalar diffusion coefficients [9]. The recent surge in the construction of physically-resolved cell models [10] responds to recognition that such colloidal physics, ranging from crowded diffusion to metabolic pathway compartmentalization, regulate cell function. In the present work, we expand our model of *Escherichia coli* to investigate how the ribosome has evolved its structure specifically to optimize its rate-limiting interactions within crowded cytoplasm (**Fig. 1A**).

**Figure 1:**
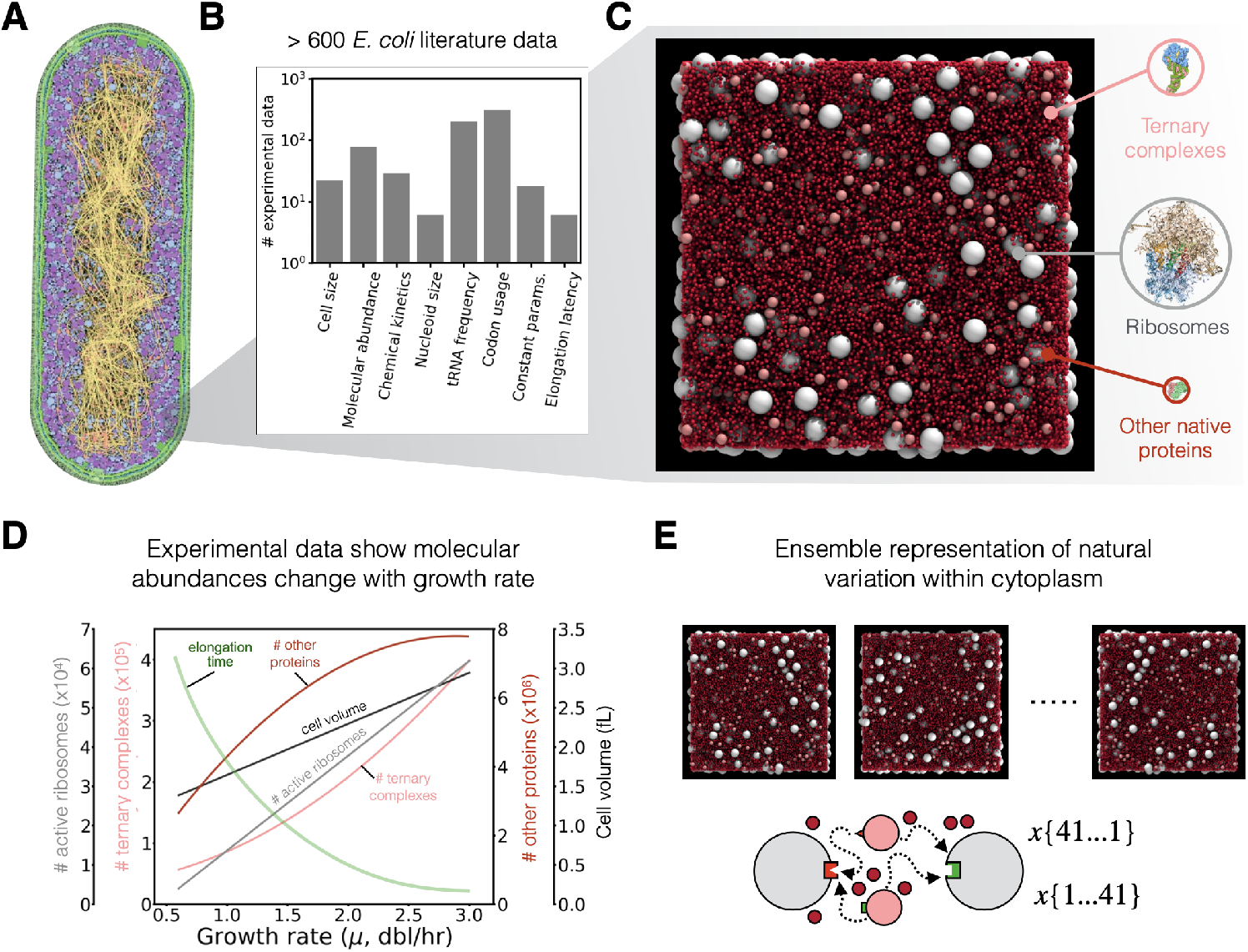
Construction of physiologically accurate *E. coli* cytoplasm in a physically-resolved computational model using extensive experimental database. **(A)** Sketch of *E. coli* [17], with permission. **(B)** Over 600 data points were compiled from *E. coli* literature to build the computational model. **(C)** Simulation snapshot of an *E. coli* translation voxel at growth rate 1 dbl/hr, containing ribosomes (white), EF-Tu GTP aa-tRNA ternary complexes (pink), and cytoplasmic proteins (red). Molecular representations adapted with permission from [13], and [17, 18] under a CC-BY-4.0 license. **(D)** Stoichiometric crowding: our model captures growth-rate dependent changes in molecular abundance and density. Figure adapted from [8], with permission. **(E)** Thousands of voxels form ensembles to represent physiological cytoplasm (adapted from [8], with permission).

One such structural feature is the ribosomal stalk protein, which interacts with translational GTPases, including elongation factors Tu (EF-Tu) and Ts (EF-Ts) [11–13]. The disordered stalk proteins, called ‘L12’ in bacteria, exist in multiple copies per ribosome, and have been shown to bind multiple GTPases simultaneously, a physical sequestration known as ‘factor pooling’ [14, 15]. Biological tests of ribosomal stalks show that L12 are essential to translation; upon knockout to fewer or no L12, elongation in *E. coli* slows dramatically or ceases altogether [12, 13]. Taken together, these biological and physical organization measurements prompt the hypothesis that ‘factor pooling’ of ternary complexes near the ribosome aids mRNA translation. Direct mechanistic interrogation of this factor pooling in bacteria is hindered by stalk flexibility, which produces L12 motions too fast to capture binding dynamics in experiments [12]. *In vivo* particle tracking experiments provide strong support that the EF-Tus are co-located near *E. coli*’s L12 stalks [16]. Understanding exactly how L12 supports ribosomal function and translation rate is important for full characterization of bacterial protein synthesis rates as well as potential drug platforms that target the ribosome. Based on experimental observations noted above and our modeling results, we hypothesize that bacterial L12 stalks’ factor-pooling function influences the A-site’s wait time for new codon recognition testing, and that the specific copy number for *E. coli* is related to its maximum growth rate.

Computational modeling provides access to cytoplasmic interactions that are still challenging to capture in experiments. To interrogate factor pooling and its influence on elongation rate, we must monitor many detailed molecular reactions and interactions at the L12, but also the cytoplasm-scale transport of ternary complexes to the ribosome. To model this process with both physical and biochemical resolution, we extend our previously developed computational model of *E. coli*, which explicitly mimics biological translation in physiological, crowded cytoplasm as it varies with growth rate [8]. Our model of the *E. coli* cytoplasm comprises physiologically accurate relative abundances of ternary complexes, ribosomes, and other cytoplasmic proteins (**Fig. 1B,C**). We represent explicitly and individually each of these molecules as they interact with each other and the surrounding cytosol. We built the model from hundreds of published experimental measurements for cell mass, cell volume, protein sizes and abundances across several growth rates, as well as *in vitro* translation elongation kinetics (**Fig. 1B** and **Table 1**). Within the model cytoplasm, molecular abundances, densities and crowding vary with growth rate, according to literature values (**Fig. 1D**) [3, 19–24]. We further represent the natural variation within cytoplasm by constructing ensembles of voxels that capture the nonuniform distribution of unique ternary complexes and nonuniform codon usage (**Fig. 1E**). The algorithm itself is not the primary focus here but rather the coupling between colloidal-scale physics and biochemistry that is key to cell function; the model is simply the tool to demonstrate this fundamental functionality. Further details of model construction and validation to *in vivo* measurements are provided in the **Methods**.

**Table 1:** Physiological data collection summary, from *E. coli* literature. Sources as shown.

| Parameter | Measured Quantity | # Growth Rates | Method | Sources | Strains |
| --- | --- | --- | --- | --- | --- |
| cell size | volume | 11 | Fluorescence microscopy | [25–27] | BW25113 and MG1655 |
| nucleoid size | volume | 3 | Light microscopy | [28] |  |
| packing | volume fraction | 6 | Calculated via equation (9-12) in [8] | [29, 30] | B/r, W1485 |
| molecular abundances | # of ribosomes | 9 | See note p of table 3 in [29] | [29] | B/r |
|  | # of elongating ribosomes | 9 | Calculated from ribosomes/cell multiplied by constant factor 0.85, as described by [29]. | [29] | B/r, W1485 |
|  | # of ternary complexes | 6 | Assumed to be equal to the number of charged tRNA per cell; from radiolabelling, ribosomal occupation measurements | [29, 30] | B/r, W1485 |
|  | # of other cytoplasmic proteins | 6 | Regression fit, on original data from Folin phenol colorimetric assay | [29] | B/r |
| molecule sizes | ribosome radii | 1 | X-ray diffraction | PDB 4V4Q from [31] | MRE600 |
|  | ternary complex radii | 1 | X-ray diffraction | PDB 1B23 from [32] | unspecified by source |
|  | cytoplasmic proteins | 1 | Averaging across single cell mass spectrometry data and dividing by average protein density. | [26] | BW25113 |
| chemical kinetics | Initial binding rate $k1, r$ | N/A | Fluorescence stopped-flow experiments and quench-flow assays. Rates adjusted from 20C to 37C using method in [33] | [3] | MRE600 |
| | Codon recognition rates $k2f, cog, k2f, nr$ | N/A | Fluorescence stopped-flow experiments and quench-flow assays. Rates adjusted from 20C to 37C using method in [33] | [4] | MRE600 |
| | GTPase activation $k3, cog, k3, nr$ | N/A | Quench-flow assays. Rates adjusted from 20C to 37C using method in [33] | [4] | MRE600 |
| | GTP hydrolysis, EF-Tu conf. change $k4, cog$ | N/A | Quench-flow assays. Rates adjusted from 20C to 37C using method in [33] | [5] | MRE600 |
| | Peptide bond formation $k5, cog$ | N/A | Quench-flow assays | [6] | MRE600 |
| | translocation $k6, cog$ | N/A | Quench-flow assays | [7] | MRE600 |
| Ternary complex frequency | relative abundance of 42 unique types | 5 | Calculated by molar ratio measurements | [30] | W1485 |
| Codon usage | frequency | 5 | Radio labeling and 2D gel electrophoresis | [30, 34, 35] | B/r, W3110 |

Using this approach, we solved the mystery of how 15-fold faster protein synthesis in faster-growing *E. coli* occurs with only 9-fold increase in ribosome abundance, showing that Brownian physics undergird the speedup and regulate translation rate [8]. Specifically, elongation quickens with growth rate due to optimized encounters between translation molecules and the ribosome in more crowded cytoplasm with more favorable stoichiometry [8]. A key point is that faster translation rate could not be explained via faster ribosome kinetics [33] nor the well-known increase in number of ribosomes. Combinatoric Brownian dynamics outside the ribosome power this increased ribosome productivity. Detailed protein/protein interactions contributed directly to cognate and non-cognate interactions and cannot be captured by simple diffusion coefficients. While the physico-chemical model thus successfully predicted *in vivo* measurements of *E. coli*’s translation speedup [3–7, 29], a quantitative gap — predicting *absolute* elongation rate — remains (**Fig. 2A**). We hypothesize that L12 stalks contribute to absolute elongation rate.

**Figure 2:**
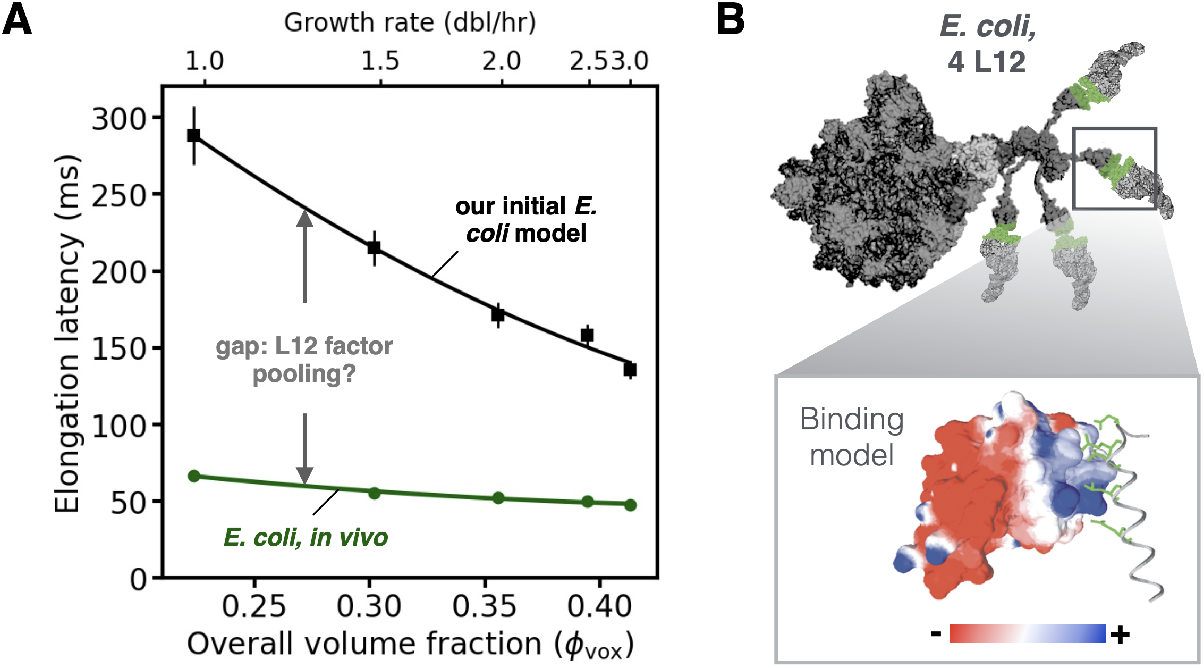
Proposed impact of L12 factor pooling on absolute elongation rate in *E. coli*. **(A)** Initial simulations of hard-sphere translation voxels (black symbols, [8]) recovered the qualitative speedup in elongation latency measured *in vivo* (green line), but left a quantitative gap (grey arrow). The *in vivo* data represent a well-established trend-line, fit through datapoints compiled from multiple independent studies [42–45, 47, 48], following the approach used in previous work [8, 48]. **(B)** Structural models of ternary complex binding to L12 ribosomal subunits in *E. coli*. (Inset) Contact scheme between the C-terminal domain of L12 subunit (red- and blue-colored surfaces) and EF-Tu helix D in the ternary complex (ribbon structure). Adapted from [12], with permission. The surface representation is colored according to charge (red, negative; blue, positive).

In this work, we expand our existing model of the growth rate dependent *E. coli* cytoplasm (**Fig. 2A**) to incorporate detailed interactions between ribosomal L12 stalks and EF-Tu (**Fig. 2B**). By explicitly modeling the transport dynamics and chemical kinetics of translation elongation, we directly interrogate factor pooling as a function of L12 copy number and cellular growth rate. Our results show that physiological EF-Tu·L12 attractions co-localize the translation machinery while maintaining high molecular mobility. This co-localization reduces wait times at the ribosomal A-site, thereby speeding protein synthesis, while promoting combinatoric sampling. We also asked whether *E. coli* could translate even faster if L12 copy number, *c*, was higher. We tested artificial conditions of *c* = 6 and *c* = 8, and identified the mechanistic tradeoff that helps explain why *c* = 4 is the optimal copy number in *E. coli*. We close with a discussion of broader implications of optimal L12 copy number across other bacterial species and implications for cellular fitness.

## RESULTS

### Modeling physiological EF-Tu·L12 interactions recapitulates factor pooling, yet avoids non-specific cytoplasmic aggregation

We modeled limited-valency binding between ribosomal L12 stalks and EF-Tu (**Fig. 2B**) with bumpy surface patches to represent the four L12 per *E. coli* ribosome and one EF-Tu per ternary complex (**Fig. 3A**). The discrete patches are spherical caps of radius 1.2 nm and are endowed in the model with attractions representing the reported binding characteristics of EF-Tu·L12 interactions [12, 36]. This binding site size is the experimentally measured width of the L12 C-terminal domain [12]. Derivation of binding parameters and detailed description of model construction are given in ***Methods***. When a ternary complex diffuses near the ribosome in our model, any contact between the two molecules outside the EF-Tu·L12 binding region is purely repulsive, reflecting the negative charge of ribosomes and ternary complexes. One to four ternary complexes can be bound to each ribosome in cytoplasm. We implemented the limited valency framework into our *E. coli* cytoplasmic voxels for a range of growth rates, monitoring positions of all ribosomes, ternary complexes, and proteins, along with reactions and transport time. As these biomolecules moved in cytoplasm via Brownian diffusion, we monitored their encounters and reactions with each other.

**Figure 3:**
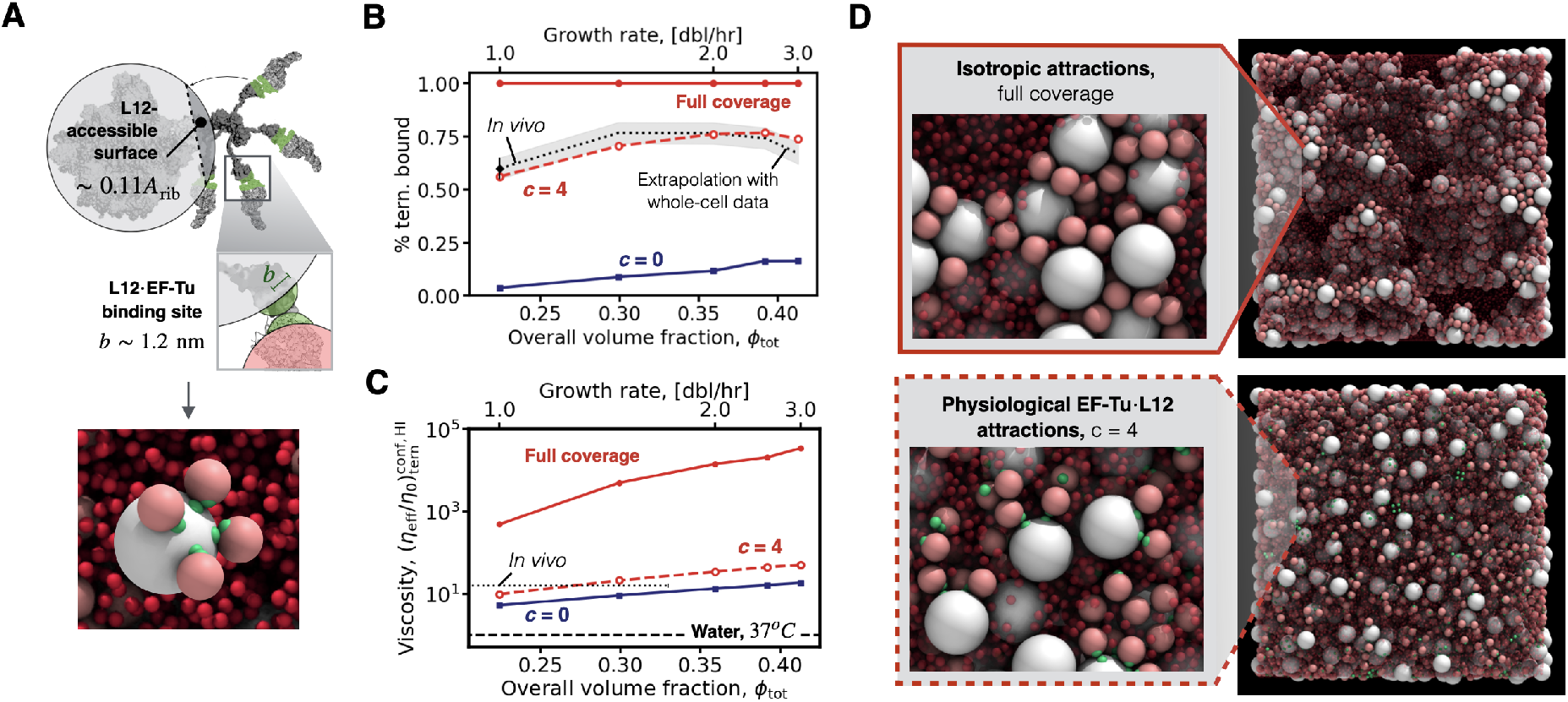
Model with natural *E. coli* L12 copy number recovers experimentally measured *in vivo* factor pooling with optimum cytoplasmic dynamics. **(A)** The physiological, limited-valency model. (Upper) Model parameterization via molecular structures, where attractions between ternary complexes (pink) and ribosomes (white) are restricted to binding sites (green), see Methods. (Lower) Simulation snapshot, present study, showing 4 ternary complexes bound to ribosomal L12 sites, embedded in dense cytoplasm. **(B)** The percent of ternary complexes bound to ribosomes (Eq. 11) and **(C)** effective cytoplasmic viscosity (Eq. 10) experienced by ternary complexes, plotted as a function of growth rate and volume fraction. The physiological model with four L12 (*c* = 4, red dashed line) recovers *in vivo* co-localization with ribosomes, while having no deleterious effect on cytoplasm viscosity. Without L12 (*c* = 0, blue solid line), ternary complexes fail to pool near ribosomes but maintain mobility. Ribosomes and ternary complexes with isotropic attractions (full coverage, solid red line) gives unphysical results. The experimentally-measured value is marked for a growth rate of 1.0 dbl/hr [16], and extrapolated to higher growth rates (dotted black line) as described in ***Supplementary Fig. S1***. The viscosity of water, *η*_0_ , at a temperature of 37C (horizontal dashed line) and the estimated *in vivo* viscosity of a protein equal in size to a ternary complex in *E. coli* (horizontal dotted line [37]) are marked. **(D)** Simulation snapshots of crowded cytoplasm voxels at a growth rate of 3 dbl/hr. (Upper) ribosomes completely covered with isotropic attraction and (Lower) physiologically accurate, limited-valence models of *c* = 4 L12. See Methods for full range of simulation conditions. Cytoplasmic proteins (red) are made translucent to improve visual clarity.

To interrogate the influence of L12 on factor pooling, we measured the fraction of ternary complexes bound to ribosomes in cytoplasm for ribosomes with no L12 stalks (*c* = 0), the physiological copy number (*c* = 4), and a very coarse-grained L12 model of a single ‘patch’ covering the entire ribosome (*c → ∞*). The results for the bound fraction (Eq. 11) are plotted in **Fig. 3B**. As expected, ribosomes without L12 stalks (*c* = 0) underpredict *in vivo* data. In the opposite extreme, a single, full-coverage patch (ribosome totally covered by attractive L12 complexes rather than negatively charged) greatly over predicts binding of ternary complexes: 100% of all cytoplasmic ternary complexes are bound to ribosomes, an unphysiological result [16]. In contrast, our model with the natural L12 copy number in *E. coli* (*c* = 4) recovers the co-localization measured *in vivo* at 1 dbl/hr [16]. *In vivo* data are available only for this single growth rate, so we extrapolated to higher growth rates by accounting for changes in molecular stoichiometry and crowding (***Supplementary Fig. S1***). These *in vivo* values match our model’s predictions of factor-pooling. Notably, the physiological model (*c* = 4) captures the non-monotonic trend of the *in vivo* data (**Fig. 3B**). As cell growth speeds up from slow to moderate rates, crowding pushes translation molecules closer together, promoting more frequent binding. As growth rate increases further, ternary complex abundance increases more than ribosome abundance [8], leaving more free ternary complexes even when ribosomal binding sites are saturated. As a result, even though crowding increases, the changed stoichiometry reduces the relative fraction of ternary complexes bound to ribosomes. The underlying microscopic picture is revealed in ***Supplementary Fig. S2-S3***, where discrete L12 sites drive the mean coordination of ribosomes toward the physiological limit of four bound ternary complexes. This static picture of co-localization suggests changes in crowding near the ribosome that could affect dynamics.

Strong diffusivity of ternary complexes should promote rapid combinatoric testing of aa-tRNAs (and thus elongation rate), but factor pooling can locally increase crowding near the ribosome, which in turn would drive down diffusivity. To study this effect, we measured the effective viscosity of cytoplasm (**Fig. 3C**), accounting for contributions from hydrodynamics and confinement [38–41] (Eq. 10, **Methods**). As expected, when there are no L12 (*c* = 0), the cytoplasm is less viscous than *in vivo* measurements, but not much lower: it approaches *in vivo* viscosity, especially at higher growth rate. But lower viscosity actually permits high biomolecule diffusivity that is actually so fast that co-localization cannot occur — made clear in **panel B**, where binding rate for *c* = 0 is much lower than *in vivo* and *c* = 4 conditions. In the opposite limit of ribosomes totally covered in L12 (*c → ∞*), the cytoplasm gels and becomes very viscous. But for *c* = 4, our model closely predicts the *in vivo* viscosity of equivalently-sized proteins in *E. coli* [37]. Simulation snapshots in (**Fig. 3D**) illustrate the frozen structure in the gelled cytoplasm that causes dynamics to arrest (top) (see also ***Supplementary Fig. S4-S6***) as well as the co-localized, mobile scenario of *c* = 4 (bottom). Combined, panels B and C suggest an optimal co-localization. Overall, the physiological L12 copy number *c* = 4 co-localizes the translation molecules that tend to promote faster recognition testing, while preserving transport dynamics that enhance combinatoric sampling essential to cognate matches.

We next sought to understand the mechanism by which this factor pooling could or would benefit recognition testing, which we hypothesize is a reduction of the time a ribosome’s A-site must wait to be re-loaded with a new ternary complex after a previous test. To facilitate rapid loading of an unoccupied A-site, co-localization duration must be comparable to codon recognition testing [3, 8]. Using experimental data from literature, we calculated this duration time previously [8] (see ***Supplementary Fig. S7***). We measured the duration of physical binding events between ternary complexes and ribosomes in our *c* = 4 L12 model, which revealed two key results. First, we found a short-duration peak from t = 0.001 to 0.01 ms driven purely by entropic forces; that is, diffusing particles naturally spend time near each other. Second, we found that EF-TU binding at L12 in our model induces longer transient binding events ranging from t = 0.01 to 1 ms, long enough for the recognition test. Data for these tests are provided in ***Supplementary Fig. S8***. These simulation results agree with *in vivo* particle tracking experiments that inferred that ternary complexes can bind to the L12, be tested, and unbind within 2 ms [16]. Overall, simulated binding at L12 co-locates ternary complexes near the ribosome long enough to re-supply the A-site after a recognition test.

### Multivalent EF-Tu·L12 interactions preload ribosomes, shorten transport time, and speed mRNA elongation

Our simulations predict that this L12-driven factor pooling directly modulates translation elongation rate in *E. coli*. To test this influence, we measured elongation latency for the physiological copy number *c* = 4 over several growth rates, and compared it to published *in vivo* data. Elongation latency, *τ*_*elong*_, includes the time a ternary complex spends reacting at the ribosome (reaction latency), diffusing in cytoplasm or pooled around an already-reacting ribosome (combined, transport latency). The wait time at the ribosome’s A-site is part of transport latency. We reported the kinetic and modeling details for this process in Methods, with additional database information in the Supplement. We discovered that physiological EF-Tu·L12 interactions (*c* = 4) speed up elongation latency by two- to three-fold compared to our original model without L12 [8] (**Fig. 4**). This decrease emerges entirely from shortened transport latency (see ***Supplementary Fig. S9***). Mechanistically, EF-Tu·L12 interactions shorten the time ribosomes must wait for a matching ternary complex to arrive at the A-site. Whereas our first model predicted the qualitative speed-up with growth rate, the current refined model improves quantitative agreement compared with *in vivo* measurements [19]. This improved prediction of absolute elongation rate supports the mechanistic picture advanced here and our hypothesis that multivalent EF-Tu·L12 interactions preload ribosomes, shorten transport time, and speed mRNA elongation.

**Figure 4:**
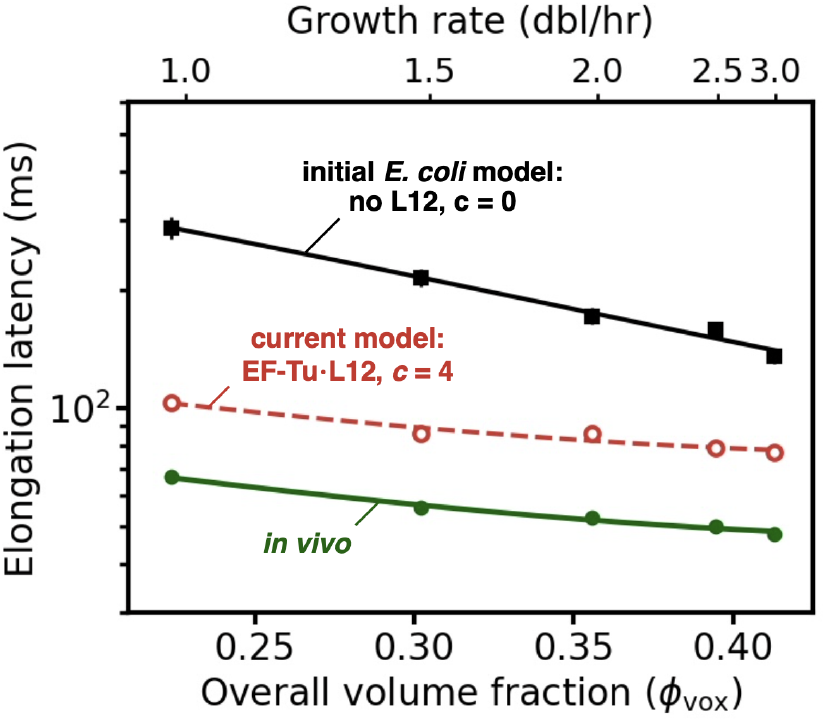
Factor pooling onto ribosomal L12 works alongside stoichiometric crowding to speed elongation. Initial simulations of translation voxels without L12 (c = 0) (black symbols, [8]) recovered the qualitative speedup in elongation latency measured *in vivo* (green line, [19, 42–48]). Physiological EF-Tu·L12 interactions (c = 4) drive two- to three-fold shorter elongation latencies (red symbols, dashed line; this work), and improve agreement with experiments. Error bars corresponding to standard error of the mean are on the order of marker size in the c = 4 case.

### More is not necessarily better: *E. coli*’s natural L12 valency optimizes its binding and sampling

Finally, we asked, “why four”? Why does *E. coli* have four L12 stalks, while other bacteria have different L12 copy number, up to eight? How does this copy number affect elongation rate? Does having higher copy number provide a faster maximum growth rate? To answer these questions, we built new ribosome models with two, six, and eight L12 binding sites, corresponding to the case when one L12 dimer is knocked out in *E. coli* (*c* = 2) [51] and physiological stoichiometries in other bacteria (*c* = 6, 8, see **Fig. 2B**) [50]. In simulations of translation voxels at 3 dbl/hr, copy numbers higher than four produced EF-Tu·L12 binding events lasting up to milliseconds (green shaded region, **Fig. 5A**). That is, more L12 reinforced factor pooling. As expected, the knock-out case, *c* = 2, dramatically reduces encounter durations and undermines the factor pooling effect. Knocking out a dimer of L12 was previously found in experiments to slow translation [51], reinforcing the connection between copy number, factor pooling, and elongation rate. Fewer than four L12 is not advantageous for *E. coli* growth.

But more is not always better. Excessive binding can come at the cost of rapid combinatoric sampling of new ternary complexes (**Fig. 5B**). We simulated translation elongation in voxels of hypothetical *E. coli* for *c* = 2, 4, 6, 8 and monitored how many unique ternary complexes, out of the entire pool, were sampled by each ribosome. High values indicate that a ribosome has many unique encounters with ternary complexes, increasing its chances for cognate interactions. As expected, the knockout case maintains the highest sampling, 87% – but encounters are over before recognition testing can begin and thus binding is ineffective. And although *c* = 6 and 8 encounters are more durable, the ribosomes sample only 55% and 45% of the combinatoric space, respectively (**Fig. 5B**). In the physiological *c* = 4 case, ribosomes are still able to sample 85% of ternary complexes on average, while also facilitating many long-duration binding events. Overall, durable binding cannot come at the cost of broad sampling — evidently *E. coli* balances them with its natural ribosomal L12 valency of *c* = 4.

We quantify this tradeoff as the product of the probabilities of the bound and freely mobile ternary complex states (**Fig. 5A-C**). Beyond the physiological copy number of four L12 per ribosome, we found that the benefits of co-localization (shorter ribosomal wait times) are wiped out by the deleterious effects of limiting ternary complex mobility (less sampling of new ribosomes) at the densest, fastest *E. coli* growth rate. That is, *c* = 4 is the optimal copy number in our simulations, as well as in live *E. coli*. In the ***Discussion***, we speculate about how a copy number of four L12 might have evolved to maximize this tradeoff and support faster growth rates more broadly.

## DISCUSSION

Cytoplasmic physico-chemical dynamics outside *E. coli*’s ribosome regulate the rate-limiting step of codon-recognition testing at its A-site [8]. In the present work, we investigated the role of *E. coli*’s L12 stalks in these dynamics. Recent studies suggest these L12 stalks pool translation factors, including EF-Tu in ternary complexes, near ribosomes [11–13, 15, 36, 52]. We hypothesized that L12 “pre-loads” the ribosome to reduce A-site wait time for new codon recognition tests. We extended our previously developed computational model, which explicitly mimics biological translation in physiological, crowded cytoplasm as it varies with growth rate [8]. Our model of *E. coli* cytoplasm includes physiologically accurate relative abundances of ternary complexes, ribosomes, and other cytoplasmic proteins (**Fig. 1C**). We represent explicitly and individually each of these molecules as they interact. We built the model from hundreds of published experimental measurements for cell mass, cell volume, protein sizes and abundances across several growth rates, as well as *in vitro* translation elongation kinetics (**Fig.1B,D** and **Table 1**) [3, 19–24]. We further represented the natural variation within cytoplasm by constructing ensembles of voxels that capture the nonuniform distribution of unique ternary complexes and nonuniform codon usage (**Fig. 1E**). For elongation kinetics within *E. coli* ribosomes, we used Rodnina’s *in vitro* data, devising an algorithm for physiological sampling [8]. Our model of A-site codon recognition testing also followed this protocol but, critically, this testing in *E. coli* commences only after a ternary complex reaches the ribosome; it must first diffuse through crowded cytoplasm, sometimes binding to non-cognate ribosomes, and waiting for testing at occupied ribosomes. Our explicit representation of physical Brownian diffusion through crowded cytoplasm, protein/protein interactions, and competition between ternary complexes revealed this search-and-match process [8]. While our initial model was the first to predict and explain the speed-up in *E. coli*’s elongation rate during faster growth, it under-predicted absolute elongation rate by as much as three fold. Here we have addressed that gap, seeking to predict the absolute elongation rate.

In the present work, we modeled L12 stalks on ribosomes and corresponding EF-Tu binding sites on ternary complexes within that growth-rate dependent *E. coli* model. The model enables us to measure transient binding dynamics at higher resolution than currently accessible by particle tracking methods [16]. We explicitly connect these binding dynamics to codon recognition testing time and elongation rate.

We first found that physiological EF-Tu·L12 binding drives durable co-localization of the translation machinery over timescales comparable to the aa-tRNA recognition test. Next, we showed with direct measurements that L12 subunits modulate protein synthesis rate by pooling ternary complexes near the ribosome. This pre-loading as much as doubles the elongation rate and substantially improved our prediction of absolute elongation rate as measured *in vivo*. Mechanistically, L12·EF-Tu binding cooperates with growth-rate dependent stoichiometric crowding to improve per-ribosome productivity by reducing A-site wait time. Our results broaden support of the recent finding that stoichiometric crowding speeds protein synthesis in faster-growing *E. coli* [8]. Beyond EF-Tu, our findings also implicate other L12-mediated interactions, including the recently identified CRISPR-Cas-associated protein Cami1 [53].

Opportunities for improvement remain. The improved but remaining quantitative gap between predicted and measured absolute elongation rate could be explained by additional physico-chemical mechanisms. For example, binding of mRNA codons to tRNA anticodons [54] could ‘pre-sort’ cognate ternary complexes near ribosomes. Modest discrepancies between *in vitro* chemical kinetic parameters and *in vivo* rates – arising, for example, from differences in solvent conditions – could also reduce the gap [8]. Future work might further refine physical interaction details to include representation of surface roughness, repulsions between negatively charged tRNA and rRNA, or the mobility of the L12 arms. Experimental data for dynamics and co-localization are limited to a single growth rate in our studied range [16]. *In vivo* data at faster *E. coli* growth rates would enable further validation of crowding-dependent behaviors in our model.

We broadened this mechanistic finding to its context in microbial selective pressure, asking “why four L12?” Bacteria exhibit natural copy numbers between four and eight (*c* = 4-8) [50]. Indeed, after surveying and collating findings from the experimental literature , we found that bacteria with *faster* growth rates tend to have *lower* L12 copy numbers (**Figure 5D** [49, 50]). To our knowledge, this correlation and apparent paradox has not been previously reported. The finding that faster-growing bacteria in general tend to have fewer L12 seems to contradict the factor pooling hypothesis. To resolve this apparent contradiction, we demonstrated one mechanism underlying the optimal L12 stoichiometry in *E. coli* (c = 4). Too many L12 effectively sequesters a local pool of ternary complexes close to each ribosome; a dissociated ternary complex is highly likely to rebind to a different L12, excluding new ternary complexes. This sequestration creates a smaller overall sampling pool for the ribosome, thus reducing favorable combinatoric sampling of new ternary complexes. In contrast, with too few L12, ribosomes fail to bind ternary complexes long enough to improve pre-loading for the A-site. The physiological *E. coli* L12 copy number, c = 4, optimizes combined durable binding and advantageous sampling of new ternary complexes.

**Figure 5:**
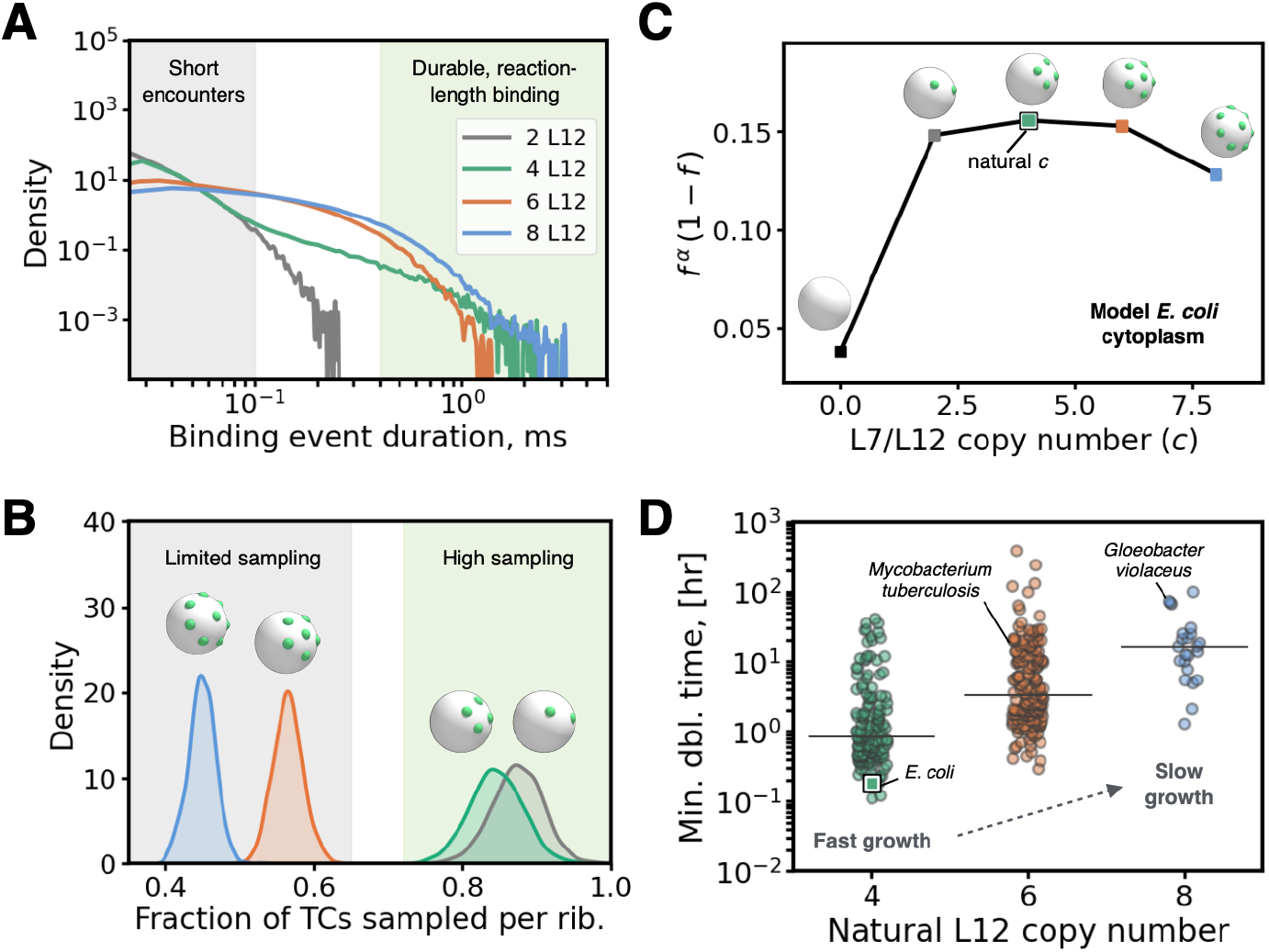
Physiological L12 stoichiometry optimizes ternary complex binding and sampling at the fastest *E. coli* growth rate. **(A)** Probability density function (shown as the kernel density estimate, KDE) of the duration of binding events for simulations with L12 copy numbers *c* = 2-8. Binding occurs when surface-to-surface distance between ternary complex and ribosome *≤* 2.1 nm. Green background highlights binding events of duration comparable to duration of initial aa-tRNA recognition test. Grey background highlights encounters too short for the recognition test to take place. **(B)** Fraction of ternary complex population sampled by each ribosome for four L12 copy numbers. Legend same as (A). **(C)** The tradeoff between bound and freely searching ternary complexes in *E. coli* translation voxels at growth rate 3 dbl/hr, as a function of L12 copy number c. Here, f is the fraction of ribosome-bound TCs and *α* = 1.71 is the ratio of Boltzmann state energies that produces *f* ^*in vivo*^ (see Methods).**D** Distributions of minimal doubling times for N = 354 bacteria [49] with known L12 copy numbers (N_4_ = 149, N_6_ = 182, N_8_ = 23, [50]). The line in each plot represents the median value. The fastest observed doubling time of *E. coli* (with c = 4 L12) is marked with a white outlined square. Distributions were compared using the Mann-Whitney U-test (p < 0.001 in all cases).

This competition suggests evolutionary selection that could favor faster or slower growth, depending on the organism. Given the hypothesis that the last bacterial common ancestor had six L12 [50], we speculate that copy numbers of c = 4 and c = 8 could have emerged to support faster and slower growth, respectively. For example, a copy number of four L12 could have evolved to optimize sampling and binding of ternary complexes under crowded conditions – that is, if the densification of cytoplasm with growth rate is a general feature of bacteria. Faster and slower growth strategies permit other survival mechanisms such as dormancy to dominate [55]. We note that cyanobacteria such as *Trichodesmium erythraeum* and *Gloeobacter violaceus* make up the majority of bacteria with eight L12, suggesting unique evolutionary pressure that favors a cytolasmic environment with enhanced binding over sampling. We encourage further work to explore the implications of this hypothesis in experiment – e.g., how L12 copy number might be engineered in bacteria to tune maximum or minimum growth rate.

## METHODS

### Modeling framework: Construction of physiological *E. coli* cytoplasm *in silico*

Rather than construct a whole-cell model of the entire *E. coli* cytoplasm, we chose to study translation elongation in voxels of cytoplasm (see discussion in ***Supplementary Information*** about computational protocols for whole-cell modeling). A voxel is a cube-shaped volume of cytosol populated with a complete set of the 42 unique ternary complexes in *E. coli*. To these voxels, we added a set of ribosomes, ternary complexes, and cytoplasmic proteins. The number of ribosomes relative to ternary complexes was set by existing literature data and ranged from four to nine per 42 ternary complexes. This number varies across growth rates, encoding variation into the construction of each voxel following [3, 8, 19–24]. These studies used to build our model were entirely from *E. coli* data, which included multiple strains as is typical [9, 56, 57], according to what is available in the literature. The modeling of L12 stalks is detailed below in paragraph entitled “Limited valency EF-Tu·-L12 interaction model”.

At a given growth rate, this approach results in one starter voxel. For example, at 0.6 db/hr, the starter voxel contains 42 ternary complexes, 4 ribosomes, and 1,970 other proteins [3, 8, 19–24]. *In vivo* data for whole *E. coli* shows that at a given growth rate, there is a natural variation in the number of cognate pairs of ribosomes and ternary complexes. This relative abundance of cognate and non-cognate pairs has been reported as relative abundances of different types of ternary complexes and frequencies of codons among mRNA in *E. coli* at different growth rates at https://doi.org/10.5281/zenodo.7200121. We constructed ensembles of translation voxels at each growth rate that together are statistically representative of these in vivo measured distributions, as follows. Using *in vivo* data, we computed the probability distributions of cognate ternary complexes at each growth rate. We then used these distributions to sample the probability that only one ternary complex matches a selected ribosome, the probability that two TCs will match that ribosome, the probability that three will match, and so on, up to the probability that all 42 ternary complexes will match the selected ribosome. We sampled all permissible configurations using reported *E. coli* codon usage and whole-cell tRNA abundances (see Table S6 and Fig. S4 at https://doi.org/10.5281/zenodo.7200121). At a growth rate of 0.6 db/hr, the chance of having 14 or more cognate TCs was vanishingly small. We thus constructed 13 additional voxels to represent the *in vivo* statistical distribution for *E. coli*, with 90 replicates per configuration. Then, for each translation voxel, one ribosome was selected at random for tracking cognate and non-cognate reactions. The speed with which the single chosen ribosome in a translation voxel finds and reacts with a cognate ternary complex provides a reliable lower bound for the bulk translation elongation rate. The bulk elongation rate corresponds to the speed with which as many peptide bonds are formed as there are ribosomes; and the speed with which a single ribosome finds and successfully reacts with a cognate ternary complex will typically be higher than the speed of as many successful reactions as ribosomes in the voxel.

### Modeling chemical reactions and elongation latencies

To simulate the coupled physics and chemistry of translation elongation, we developed expanded functional modules for LAMMPS [58] to model changes in interparticle interactions based on distance, time, and particle type. To represent codon-anticodon testing, we modified our limited-valency interaction model to include internal sites within each ribosome and ternary complex, representing the A-site codon and tRNA anti-codon, respectively (see ***Supplementary Fig. S13*** ).

In simulation, we allowed particles to diffuse and interact with both entropic hard sphere repulsion and EF-Tu·L12 attractions. To simulate the initial aa-tRNA recognition test, we used a distance-based rule to change the pairwise interactions between ribosomes and ternary complexes that are in close physical proximity. Namely, if the center-to-center distance between L12 and EF-Tu sites is less than 1.05 times their hard-sphere contact distance (i.e., 2.52 nm), we considered their respective ribosome and ternary complex as having begun reacting. We note that molecules can only be involved in one reaction at a time. To simulate the entry of a captured ternary complex into the ribosome, we turned off the hard sphere repulsion between reacting pairs, and turn on a 16 *k*_B_*T* attraction between their internal A-site codons and tRNA anti-codons (see ***Supplementary Fig. S13***). The duration of each reaction was drawn from a latency distribution derived from experimental kinetic rates [8], depending on if the codon and anticodon are cognates or non-cognates (see ***Supplementary Fig. S7***). We accounted for the presence of near-cognate ternary complexes by randomly scaling non-cognate reaction durations by 3.3-fold with a 20% probability, as in [8]. If the reaction was not successful, the simulation was allowed to proceed for the sampled reaction duration, before ternary complex release was imposed by a soft repulsion between the reacting pair. Once the center-to-center distance was above a distance threshold (12.2 nm), both particles in the simulation were returned to the non-reacting pool and allowed to start another reaction upon their next encounter. When a successful cognate match was found, the simulation was terminated.

The previous reaction framework [8] scaled physiological kinetic rates by 600-fold, which enabled computational tractability while maintaining a well-mixed distribution of particles within simulation. Since physical binding events in our limited-valency model last ten times longer on average than in non-attractive voxels, we scaled kinetic rates here by only 60-fold. For more detail, see a discussion of this speedup in the ***Supplementary Information*** section “Acceleration of translation voxel simulations to reduce run time and cost.”

For each growth rate, we calculated a reaction latency 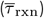, defined as the time the cognate ternary complex spends in reactions; an elongation latency 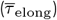, defined as the total time the matching ribosome spends unbound or in reactions; and a transport latency 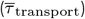, calculated as the difference between the elongation and reaction latency. Each was averaged across *j* = 90 replicates, weighted by the probability of a given number of cognate ternary complexes, up to *i* = 13 per voxel:

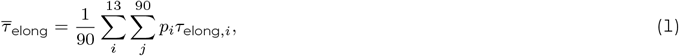

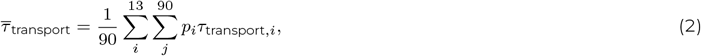

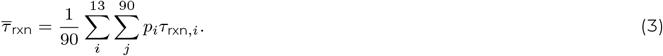

### Modeling intermolecular interactions

We developed a framework in which pairs of molecules *i* and *j* can interact in translation voxels via entropic or attractive components, defined by the potential:

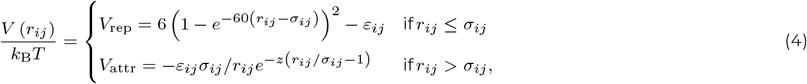

where *r*_*ij*_ is the magnitude of the unit vector 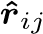 pointing from the center of particle *j* to the center of particle *i, k*_B_ is the Boltzmann constant, and *T* is the absolute temperature. Entropic exclusion was enforced via a Morse potential, *V*_rep_, previously demonstrated to recover hard-sphere behavior [59]. The hard-sphere contact distance *σ*_*ij*_ is defined as the sum of particle radii, *a*_*i*_ and *a*_*j*_ . Previous structural homology-based and kinetic experiments suggest that the EF-Tu·L12 interaction consists of a hydrophobic patch flanked by two salt bridges [12, 36]. To capture the exponential character of such interactions [60, 61], we represented the attraction between ternary complexes and ribosomes via a Yukawa potential, *V*_attr_. The attraction range of hydrophobic interactions and salt bridges (approximately 0.4 nm [61, 62]) corresponds with a decay rate constant *z* = 10*/σ*_*ij*_ in Eq. 4. The characteristic attraction strength, *ε*_*ij*_ , sets the depth of the potential well. We estimated the *in vivo* attraction strength to be 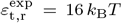, assuming each salt bridge contributes 3 kcal*/*mol (4.9 *k*_B_*T* at 37^*o*^*C*, [63]) and the hydrophobic patch contributes 4 kcal*/*mol (6.5 *k*_B_*T* at 37^*o*^*C*). To calculate the hydrophobic contribution, we assumed that a 18 cal mol^−1−2^ free energy density [64] is distributed over a hemisphere of diameter 11.8 , the size of the hydrophobic region of the L12 C-terminal domain (PDB code ICTF [65]). We also investigated weaker isotropic attraction strengths of *ε*_t,r_ = 3 *k*_B_*T* and 6 *k*_B_*T* , which have been demonstrated to drive formation of phase-separated droplets [66] and arrested gel networks [67], respectively. Interactions between other types of molecules were modeled as purely repulsive at contact and otherwise did not interact, i.e., *ε*_*ij*≠t,r_ = 0.

### Limited valency EF-Tu·-L12 interaction model

To build a limited-valency interaction model, we first calculated the ribosome surface area accessible to the highly mobile L12 subunits. Using the structural model from [13], we projected the range of movement of all four L12 subunits onto a ribosome body of size *a*_rib_ = 13 nm (**Fig. 3A**). The resulting spherical cap with base radius *h* = 0.61 *a*_rib_ has a surface area *A*_cap_ = 220.4 nm^2^, approximately 11% of the total ribosome surface area *A*_rib_ = 2123.7 nm^2^. We embedded four spheres representing the L12 subunits at evenly spaced intervals around the boundary of this accessible surface area. These spheres were inset to expose small binding site caps with base radii *b* = 1.2 nm, the measured size of the L12 C-terminal domain (PDB code *ICTF* [65]). All measurements of structural models in this work were conducted using UCSF ChimeraX [68]. We embedded an identical inset sphere in the surface of each ternary complex to represent the equivalent binding site on EF-Tu (**Fig. 3A**). In the limited-valency model, the Yukawa attraction *V*_attr_ (Eq. 4) is only felt between these inset binding sites representing the EF-Tu and L12 domains.

To interrogate a range of L12 copy numbers, *c*, we modeled ribosomes with two, six, and eight binding sites of identical size to the previously described *c* = 4 model. For *c* = 2, we inset the binding sites on opposite sides of the L12 accessible surface area (**Fig. 5A, C**). For *c* = 6 and *c* = 8, we inset one binding site at the center and five and seven binding sites, respectively, equally spaced around the circumference of the L12 accessible surface area. When *c ≥* 6, we expanded the accessible surface area to maintain the same binding site spacing as in the *c* = 4 case to prevent a single ternary complex from binding to multiple L12 at a time.

### Dynamic simulation method

Molecules were immersed in an implicit Newtonian solvent with viscosity *η* = 0.6913 mPa*/*s and density *ρ* = 0.99336 g*/*cm^3^, chosen to represent water at 37^*o*^ C. The small size of translation molecules set a vanishingly small Reynolds number, *Re* = *ρUa*_*i*_*/η*, where *U* is the characteristic particle velocity. Fluid motion in this simulation was thus governed by Stokes’ equations, whereas particle motion was governed by the Langevin equation [69]:

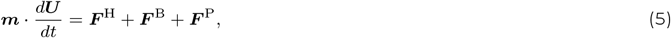

where ***m*** is the mass tensor and ***U*** is the particle velocity. ***F*** ^H^ is the hydrodynamic force on each particle *i* given by Stokes drag law 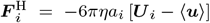, where ⟨***u***⟩ is the bulk fluid velocity, which is zero for a quiescent prokaryotic cytoplasm. The stochastic Brownian force ***F*** ^B^ emerges from thermal solvent fluctuations and obeys Gaussian statistics:

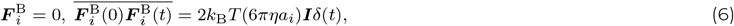

where overbars indicate averages over long times compared to the solvent timescale, ***I*** is the identity tensor, and *δ*(*t*) is the Dirac delta function. The deterministic interparticle force ***F*** ^P^ is the gradient of the interaction potential *V*_*ij*_ between a particle pair (Eq. 4):

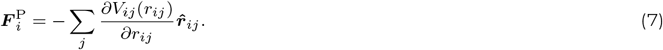

We instantiated the initial distribution of finite sized molecules in translation voxels using PACKMOL [70]. In LAMMPS, Eq. 5 is integrated forward in time using the velocity-Verlet method [71] and a Langevin thermostat [72] to solve for particle trajectories. We chose a sufficiently small discretization timestep, Δ*t* = 13 ps, and damping factor, *d* = 0.01, to recover the inertia-less (overdamped) physics of the Stokes flow regime [59]. In the limited-valency model, each ribosome and ternary complex with inset binding sites was treated as a rigid body using the ‘fix rigid’ command in LAMMPS. Simulations were evolved for 1.3 ms, with full particle trajectories output every 130 ns. Visualizations of simulations were created using Visual Molecular Dynamics (VMD, [73]).

### Dynamics and rheology

We monitored dynamics by tracking particle positions throughout the simulation and computing their mean-square displacement (MSD) over time

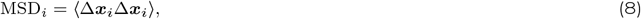

where Δ***x*** = ***x***(*t*) − ***x***(0) is the change in particle position from the initial time to the current time *t*, and *i* is the particular molecular species (ribosome, ternary complex, or small native protein). Please see ***Supplementary Fig. S10-S11***. Angle brackets denote an ensemble average over all molecules of that species. We extracted the long-time self-diffusivity, 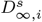 for each species from the slope of the MSD over long times

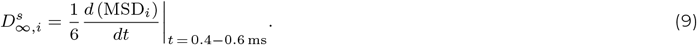

Cellular confinement and the presence of hydrodynamic interactions are expected to slow diffusion *in vivo*. Thus, to compare dynamics in our freely-draining, unconfined model to experimental measurements, we scaled 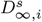 by a volume fraction dependent factor which accounts for both confinement and hydrodynamic effects. We derived this scaling factor using analytical theory [74, 75] and results from confined Stokesian and Brownian Dynamics simulations [39, 41]. Details can be found in the ***Supplementary Information, Fig. S12***.

We used this molecular diffusivity to calculate the effective cytoplasmic viscosity experienced by ternary complexes *η*_eff,tern_, given by the Stokes-Einstein relation [76, 77], which is a reasonable approximation in concentrated suspensions if the produced viscosity is interpreted as a microscopic rather than a bulk viscosity [78]:

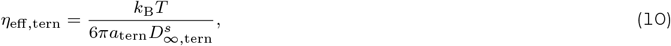

Here, 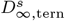 is the scaled long-time self-diffusivity, accounting for hydrodynamic and confinement effects (***Supplementary Fig. S12***). We normalized this effective viscosity by that of water at 37^*o*^ *C, η*_0_ = 0.6913 cP, to capture the entropic contribution due to finite molecular size.

### Binding metrics

To quantify co-localization of the translation machinery, we calculated the fraction of ribosome-bound ternary complexes, *f* , in each voxel:

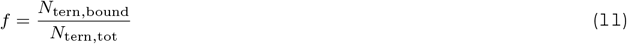

where *N*_tern,tot_ is the total number and *N*_tern,bound_ is the ribosome-bound number of ternary complexes. A ternary complex, *i*, was considered bound to a ribosome, *j*, if their center-to-center separation *r*_*ij*_ was less than the maximum bond distance *r*_cut_ = 21 nm (for details, see ***Supplementary Fig. S8A***). This bound fraction was estimated *in vivo* by Mustafi and Weisshaar to be *f*_exp_ = 0.6 at a growth rate of 1 dbl/hr [16]. In ***Supplementary Fig. S1***, we describe how we extrapolated this experimental measurement to higher growth rates by accounting for changes in molecular stoichiometry and crowding.

### Binding times and sampling rates

We quantified the duration of individual binding events by computing the time that ternary complexes spend within the maximum bond distance *r*_cut_ of a ribosome. According to this binding definition, we tracked the state of each ternary complex as ‘bound’ or ‘unbound’ over the 1.3 ms simulation evolution. If a ternary complex departed a bound state, the duration of that event was recorded in the distribution of binding times. The identity of that ternary complex was also recorded as having been ‘sampled’ by that ribosome. At the end of the analysis window, we computed the number of unique ternary complexes that each ribosome sampled, normalized by the maximum number sampled in the *c* = 2 case. We visualized the probability distributions of binding event durations and sampling fractions using the Python Seaborn package for kernel density estimates.

We quantified the tradeoff between co-localization and mobility by the product of the fractions of bound, *f* , and freely diffusing ternary complexes, (1 − *f* ). Without scaling, this product would be maximized at *f* = 0.5, whereas *f* ^*in vivo*^ = 0.67 at a growth rate of 3 dbl/hr (**Fig. 5C**). To account for this favoring of the bound state *in vivo*, we raised the bound fraction *f* to a power *α* = 1.71, corresponding to the difference in energies (*ε*_free_ − *ε*_bound_) in a two-state Boltzmann distribution [79]:

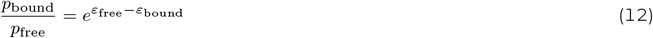

where *p*_bound_ = *f* ^*in vivo*^ and *p*_free_ = 1 − *f* ^*in vivo*^.

### Prokaryotic L12 Copy Number

L12 copy number, *c*, has been quantified for more than 1, 200 bacteria, mitochondria, and chloroplasts by computational analysis and mass spectrometry [50]. We focused on a subset of *N* = 354 bacteria across five phyla for which minimal doubling times are also known (Table S2), gathered from the ‘condensed_traits_NCBI.csv’ supplementary file of [80] (https://doi.org/10.6084/m9.figshare.12221732). We separated this subset into three populations of size *N*_*c*_ according to L12 copy number, where *N*_4_ = 149, *N*_6_ = 182, and *N*_8_ = 23. We then visualized the probability distribution of minimal doubling times for each population using the Python Seaborn package for kernel density estimates. We compared the means of the distributions via the Mann-Whitney *U* -test in SciPy, with *p <* 0.001 for all cases.

## DATA AVAILABILITY

All study data and code for analysis are available on Github (https://github.com/hofmannj-git/Elongation_EF-Tu_L7L12.git). Simulation code can be reproduced by the reader utilizing the open-source platform LAMMPS [58]. Discussion regarding our particular modifications is available upon request.

## ACKNOWLEDGMENTS

The authors thank Drew Endy, Akshay Maheshwari, and Emma Gonzalez for many fruitful discussions and insights regarding the translation voxel model. This work was supported by the National Science Foundation Graduate Research Fellowships under Grant No. 1656518 for J.L.H. and A.M.S., Stanford Graduate Fellowships for A.M.S and T.S.Y., and an ARCS Foundation Award for J.L.H.. Computing was performed on the Sherlock cluster at the Stanford Research Computing Center (SRCC) and on Stampede2 at the Texas Advanced Computing Center (TACC). SRCC resources were provided by Stanford University, while resources at TACC were supported by the National Science Foundation’s Extreme Science and Engineering Discovery Environment (XSEDE) Research Award No. CTS130035 and Advanced Cyberinfrastructure Coordination Ecosystem: Services & Support (ACCESS) Research Award No. CHM230007.

## AUTHOR CONTRIBUTIONS

J.L.H. and R.N.Z. designed research; J.L.H. and T.S.Y. performed research; J.L.H., T.S.Y., and A.M.S. analyzed data; and J.L.H., T.S.Y., A.M.S., and R.N.Z. wrote the paper.

## SI APPENDIX

### Supplementary methods

#### Rescaling of diffusivity: confinement and hydrodynamics

To compare the long-time self-diffusivities 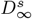 calculated in our simulations to those from *in vivo* particle tracking experiments [37], we derived a volume fraction dependent scaling factor to account for the effects of hydrodynamic interactions (HI) and confinement. The long-time self-diffusivity, 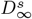, of a suspension of purely-repulsive hard spheres can be expressed [75] as:

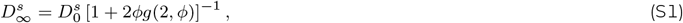

where 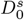 is the short-time self-diffusivity, *ϕ* is the volume fraction, and *g*(2, *ϕ*) is the value of the radial distribution function at contact for a given volume fraction *ϕ*. Our simulations represent the unbound freely-draining limit in which 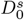 is simply *D*_0_, the Stokes-Einstein equation of a single colloid diffusing in a continuum liquid [76, 77]. Thus,

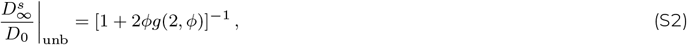

holds in the unbound, freely-draining limit (solid black line, **Fig. S12A**). The analytical form of 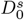 calculated by Beenakker and Mazur in 1984 [74] gives:

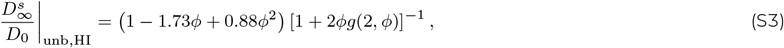

for the unbound case with full HI (solid red line, **Fig. S12A**). Analogous analytical forms of 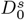 do not yet exist for confined suspensions, so we turned to results from dynamic simulations. The confined 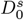 are calculated from confined Brownian (dashed black line, **Fig. S12A**; freely-draining limit [41]) and Stokesian Dynamics simulations (dashed red line, **Fig. S12A**; with HI [39]), averaged over all positions in a spherical cavity with a confinement ratio of *λ*_c_ = *a*_p_*/a*_cav_, where *a*_p_ and *a*_cav_ are the radii of the particle and cavity, respectively. In all cases, *g*(2, *ϕ*) was calculated via Percus-Yevick theory, as described in the subsequent section.

The volume fraction dependent scaling factor, *s*, is determined by the ratio of 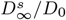 for the unbound case and confined case with HI. The scaling factor increases with volume fraction (**Fig. S12A**), as crowding compounds the slow-down in diffusion from both hydrodynamic interactions and confinement. All instances of 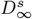 in our work are divided by the value of *s*(*ϕ*), fit to a second-order polynomial in Python (**Fig. S12B**).

#### Theory for the bidisperse radial distribution function

We computed the radial distribution function *g*(*r*) in translation voxels without small cytoplasmic proteins using Percus-Yevick (PY) theory [81]. We calculated *g*(*r*) for bidisperse voxels using the code provided in [82] to numerically evaluate PY theory for hard sphere mixtures [83, 84] representing the size and abundance of ternary complexes and ribosomes across *E. coli* growth rates (Table S1). We examined two cases. First, we computed *g*(*r*) when the total voxel volume *V*_tot_ is the same as in the tridisperse case, i.e., if crowders are simply removed (solid line, **Fig. S14B**). Second, we computed *g*(*r*) when the total volume fraction *ϕ*_tot_ matches the tridisperse case, i.e., if crowders are removed and the voxel volume is reduced to recover the lost particle density (dashed line, **Fig. S14B**). To estimate the strength of this apparent depletion interaction, we ran dynamic simulations of bidisperse voxels across all *E. coli* growth rates, with volumes *V*_tot_ matching the tridisperse case. We included a *ε*_t,r_ = 1 *k*_B_*T* attraction between ternary complexes and ribosomes (of the form in Eq. 4 and calculated *g*(*r*) as described above via Eq. S5.

#### Binding affinity

We calculated the apparent binding affinity of the interaction between ribosomes and ternary complexes, as defined in [85]:

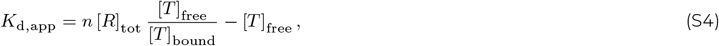

x where we assume each ribosome has *n* independent sites to which ternary complexes can bind. [*R*]_tot_ is the total concentration of ribosomes, and [*T* ]_free_ and [*T* ]_bound_ are the concentrations of free and bound ternary complexes, respectively. Ribosome-bound ternary complexes are defined as in Eq. 11, i.e., if they are found within *r*_cut_ of a ribosome. For the case with isotropic attractions, the number of independent binding sites *n* is set by the number of ternary complexes that can fit around a ribosome at maximum packing. With a relative particle size ratio of *a*_tern_*/a*_rib_ = 0.45, the isotropic number of binding sites is thus *n*_iso_ = 32 [86]. For the limited-valency attraction case, we set the number of binding sites *n*_lim_ = 5 to reflect the maximum measured coordination number of ribosomes, *N*_c_ (see **Fig. S2, Fig. S16B, Fig. S17B**)

#### Suspension Microstructure

We calculated the radial distribution function, *g*(*r*), to characterize pair-level correlations between ternary complexes and ribosomes, defined according to [87, 88]

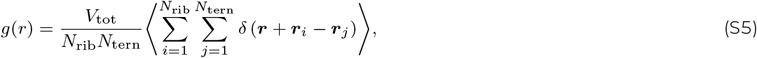

where *V*_tot_ = (*l*_vox_)^3^ is the total voxel volume and the function 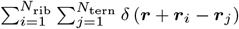 is the microscopic particle density at a given position ***r***, calculated in translation voxels with separations ***r***_*i*_ ***r***_*j*_ for all *i* ribosome and *j* ternary complex pairs. The radial distribution function is normalized to reflect departures from a uniform distribution, *g*(*r*) = 1 that arise from finite size and other effects. Please see **Fig. S17A**.

We calculated the static structure factor *S*(***q***), the wave-space equivalent of *g*(*r*) that is typically obtained in scattering experiments [67, 87] according to:

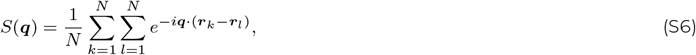

where ***q*** is the wavevector normalized by the ribosome size *a*_rib_, *N* is the number of particles, and ***r***_*k*_ − ***r***_*l*_ is the separation for a particle pair (*k, l*). *S*(***q***) is radially averaged to give *S*(*q*), where *q* = |***q***| and corresponds to a real space length of *l* = 2*π/q*. We were interested in quantifying long-range correlations throughout the translation voxel and thus calculated *S*(*q*) for all particles *N* = *N*_rib_ + *N*_tern_ + *N*_prot_ in the ensemble.

#### Coordination numbers

A ternary complex, *i*, was considered ‘coordinated’ with or bound to a ribosome, *j*, if their center-to-center separation *r*_*ij*_ is less than the maximum bond distance *r*_cut_ = 21 nm. This value corresponds to a separation of 2.1 nm beyond hard-sphere contact, which includes the radii of the binding site protrusions and the width of the attractive well. According to this definition, we calculated the number of ternary complexes to which each ribosome is bound – its coordination number, *N*_c_. We computed the distribution of ribosome coordination numbers for voxels across all growth rates and attraction strengths (**Fig. S3, S2**). We quantified the duration of individual binding events by computing the time that ternary complexes spent within the maximum bond distance *r*_cut_ of a ribosome (**Fig. S8A**).

#### Extrapolation of *in vivo* co-localization to faster growth rates

The fraction of ribosome-bound ternary complexes, *f* , has been measured *in vivo* at only one growth rate, *f* ^exp^(1 dbl*/*hr) = 0.6. We extrapolated this *in vivo* measurement to faster cellular growth rates (**Fig. S1**) by calculating the expected change due to crowding (+Δ*f*_crowd_) and stoichiometry (+Δ*f*_stoich_). We defined the change in co-localization due to purely entropic forces as:

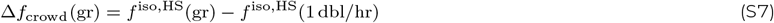

or the change in *f* with growth rate (gr) in the baseline hard sphere simulations (Fig. 3B, *ε*_t,r_ = 0 *k*_B_*T* . We define the change in co-localization due to stoichiometry as:

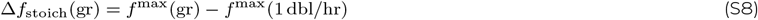

where *f* ^max^(gr) = 4*N*_rib_(gr)*/N*_tern_(gr) is the maximum possible *f* , assuming that each ribosome is bound to exactly four ternary complexes. *N*_rib_(gr) and *N*_tern_(gr) were extracted and compiled from whole-cell measurements by [8]. We summed these two contributions (+Δ*f*_tot_) to obtain the extrapolated *in vivo* bound fraction:

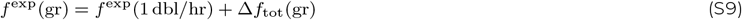

#### Translation voxel model description

The following subsections contain detailed descriptions of the translation voxel model in *E. coli* and are reproduced with permission from [8].

#### Representing crowding and stoichiometry at different growth conditions

We developed computational representations of translation voxels by analyzing the abundances and sizes of molecules comprising *E. coli* cytoplasm in relation to overall cell volume and mass. Where needed, we inferred abundances across growth rates by fitting polynomials to reported measurements (described below). This procedure allows us to accurately represent the *volume fraction*, which is the relevant metric of crowding that sets macromolecular dynamics in Stokes flow, of the cytoplasm at different growth conditions.

#### Calculation of biomolecular abundances in cells

We computed the average abundances of ribosomes (*N*_rib_) and ternary complexes (*N*_tern_ in single cells across physiological growth rates using existing data from literature (Table S1 in [8]). To compute the abundance of proteins surrounding ribosomes and ternary complexes, we first calculated the total dry mass of proteins in cytoplasm, *M*_cytoplasm,prot_. Protein mass encompasses the mass of all biomolecules in cytoplasm other than ribosomes, ternary complexes, mRNA, and DNA:

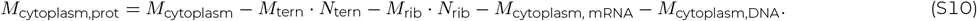

The masses of ternary complexes (*M*_tern_) and ribosomes (*M*_rib_) were specified using known molecular structures and the total mass of mRNA, DNA, and cytoplasm (*M*_cytoplasm,mRNA_, *M*_cytoplasm,DNA_,and *M*_cytoplasm_, respectively) were taken from literature (Table S1 and S4 in [8]). We then calculated the average effective spherical radius of proteins 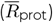, as well as the mass of resulting average-sized proteins 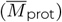, using single-cell *E. coli* mass spectrometry data and average protein density (*ρ*_prot_):

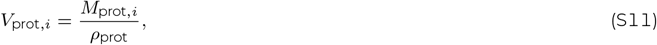

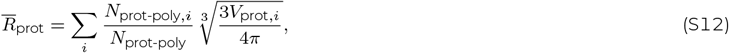

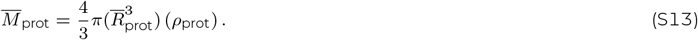

We obtained the mass and abundances of each protein in the *E. coli* cytoplasm (*M*_prot,*i*_ and *N*_prot-poly,*i*_, respectively), as well as the total number of proteins (*N*_prot-poly_), from the mass spectrometry measurements of Heinemann and colleagues [26]. We also computed the volume of each protein in the *E. coli* cytoplasm (*V*_prot,*i*_) using these measurements. We then determined the abundance of proteins in a cell (Fig. S19) as the ratio of total mass occupied by proteins in cytoplasm (Eq. S10) and the mass of average-sized proteins,

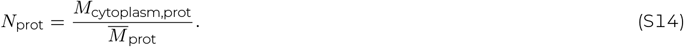

We note that the size polydispersity of proteins is weak – meaning they do not deviate much from the average size (Fig. S21). As a result, we expect negligible change to ternary complex dynamics and overall translation rates, justifying our representation of protein crowding with an average size.

#### Acceleration of translation voxel simulations to reduce run time and cost

Simulations of translation voxels without L12 were originally forecast to cost US $6 million with the longest simulations taking *∼*36 years, making them intractable. To achieve feasible costs and run times, we previously implemented a procedure for accelerating the simulations without L12 by *∼*600-fold, reducing costs and run times as detailed above. In our acceleration procedure, kinetic rates of unbinding and codon recognition (i.e., the possible exits to the initially bound state) were increased 600-fold during simulations (*k*_1_ = 717 s^−1^ to 430,200 s^−1^ and *k*_2f_ = 1,474 s^−1^ to 884,400 s^−1^ ). Simulations were run until completion following a successful match between a cognate ternary complex and matching ribosome. Subsequently, during post-simulation analysis, the time spent by ternary complexes in the initially bound state was rescaled to be 600-fold longer, and rescaled times were used to compute reaction, transport, and elongation latencies. Our estimates of overall reaction, transport, and elongation latencies were not sensitive to this scaling procedure at the *∼*600-fold acceleration used (Fig. S5 at https://doi.org/10.5281/zenodo.7200121) for voxels without EF-Tu · L12 attractions. This insensitivity is a result of unbinding kinetics remaining slow enough that, for a certain range of kinetic scaling, ternary complexes mix within the translation voxel between unbinding events to a sufficiently similar extent.

As described in the ***Methods***, we scaled kinetic rates by only 60-fold in this work, given ten-fold longer physical binding events in simulations with EF-Tu·L12 attractions and to maintain this voxel mixing assumption.

### Supplementary results

#### Static structure factor

We calculated the static structure factor, *S*(*q*), in translation voxels across all *E. coli* growth rates and isotropic attraction strengths studied in this work (**Fig. S5**). In the hard sphere limit (*ε*_t,r_ = 0 *k*_B_*T* ), we found that cytoplasmic polydispersity prevents the emergence of strong structural correlations at any wavevector *q* or real space length *l* = 2*π/q*. The probability of long-range correlations at small wavevectors is only slightly increased by weak isotropic attractions (*ε*_t,r_ = 3 *k*_B_*T* ), suggesting the formation of transient low-order clusters. A single peak emerges for the *ε*_t,r_ = 6 *k*_B_*T* attractive case, characteristic of a bicontinuous gel created by arrested phase separation. The peak shifts to higher wavevectors as crowding increases, corresponding with formation of thinner and shorter strands. In the *ε*_t,r_ = 16 *k*_B_*T* attractive case, there is an abundance of correlations at longer length-scales (shorter wavevectors), but the stringier structure is not as well characterized by a single length-scale since ongoing phase separation is slowed by strong bonding [67].

#### Effective cytoplasmic viscosity

We calculated the effective cytoplasmic viscosity experienced by ternary complexes as a function of cellular growth rate, accounting for the impacts of hydrodynamic interactions and cellular confinement (see ***Methods*** above). Without attractions (*ε*_t,r_ = 0 *k*_B_*T* ), the effective viscosity exhibits the expected increase with volume fraction, but underpredicts by two- to three-fold the *in vivo* estimate for equivalently-sized proteins in *E. coli* [37] (**Fig. S15, S18**). Transient cluster formation in the case of weak isotropic attractions (*ε*_t,r_ = 3 *k*_B_*T* ) drives an approximately 5% increase in effective viscosity, whereas stronger isotropic attractions over-predict *in vivo* viscosity by up to five-fold (*ε*_t,r_ = 6 *k*_B_*T* ) and up to two orders of magnitude (*ε*_t,r_ = 16 *k*_B_*T* ). In all cases, viscosity increases with cell growth rate as the cytoplasm densifies.

## Supplementary tables

**Table S1:** Growth-rate dependent simulation parameters in scaled-up translation voxels of *E. coli* cytoplasm

| Doublings/hr | $\mu = 1.0$ | $\mu = 1.5$ | $\mu = 2.0$ | $\mu = 2.5$ | $\mu = 3.0$ |
| --- | --- | --- | --- | --- | --- |
| $N_{\text{rib}}$ | 659 | 1,005 | 1,266 | 1,426 | 1,524 |
| $N_{\text{tern}}$ | 3,528 | 4,662 | 6,090 | 7,476 | 8,820 |
| $N_{\text{prot}}$ | 176,064 | 198,801 | 205,610 | 194,020 | 172,200 |
| $l_{\text{vox}}$ | 405.0 nm | 404.4 nm | 403.7 nm | 404.3 nm | 404.0 nm |
| $\phi_{\text{tot}}$ | 0.224 | 0.299 | 0.359 | 0.392 | 0.413 |

## Supplementary figures

**Figure S1:**
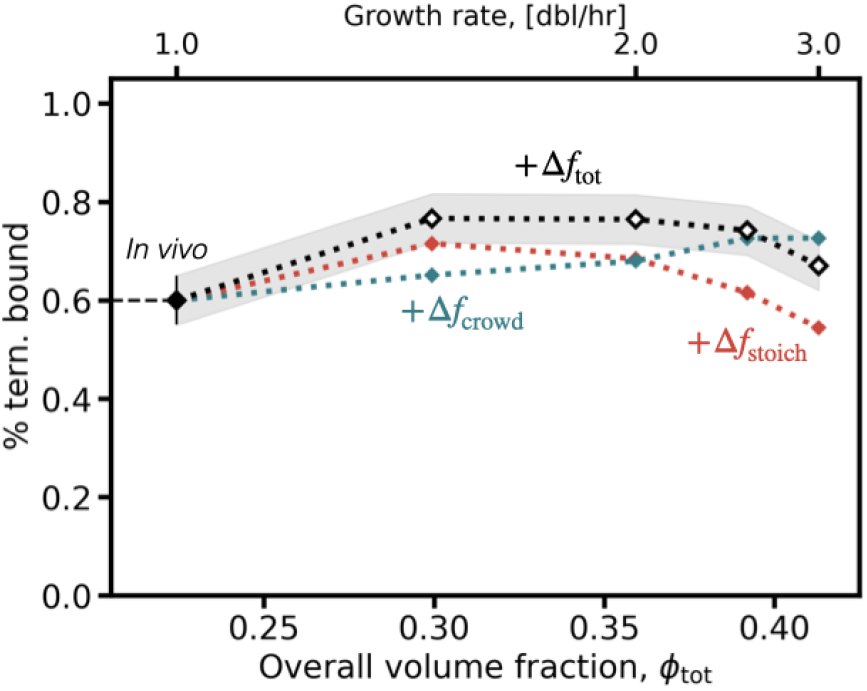
*In vivo* co-localization of ternary complexes and ribosomes is projected to increase non-monotonically as a function of growth rate. Extrapolation of the measured *in vivo* fraction of ribosome-bound ternary complexes, *f* , at 1 dbl/hour (black solid diamond) to higher growth rates (see Supplementary Methods ). The change in co-localization due to entropic forces (+Δ*f*_crowd_, teal) and changing relative stoichiometry (+Δ*f*_stoich_, orange), assuming each L12 is bound to one ternary complex. These two contributions are summed to obtain the extrapolated *in vivo* prediction to higher growth rates (+Δ*f*_tot_, black), including errors carried forward from the experimental measurement at 1 dbl/hour.

**Figure S2:**
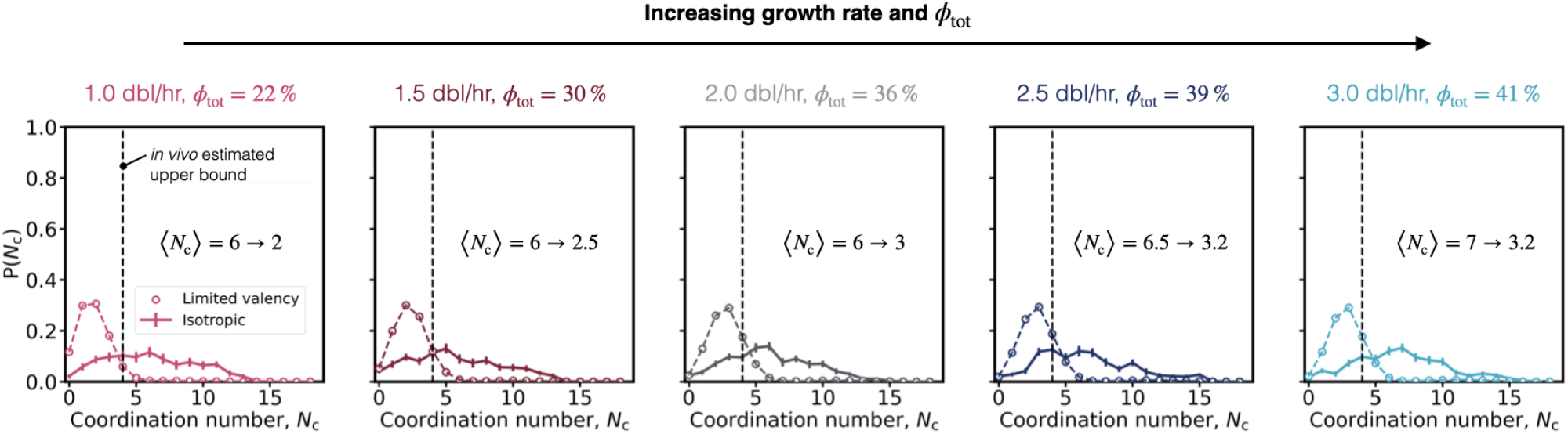
Limited valency attractions drive biofidelic co-localization of translation molecules, without higher-order microstructure. Coordination number distributions highlight an increase in the number of ternary complexes bound to each ribosome, both with increasing growth rate (red to grey to blue) and interaction valency (dashed versus solid lines) for attractions of strength *ε*_t,r_ = 16 *k*_B_*T* . The estimated maximum ribosome occupancy from experiments [16] is four ternary complexes per ribosome, marked by a vertical dashed line. A reduction in the average number of ternary complex contacts, ⟨*N*_c_⟩ , felt by ribosomes is noted for the isotropic case relative to the limited valency case, at each growth rate (left to right).

**Figure S3:**
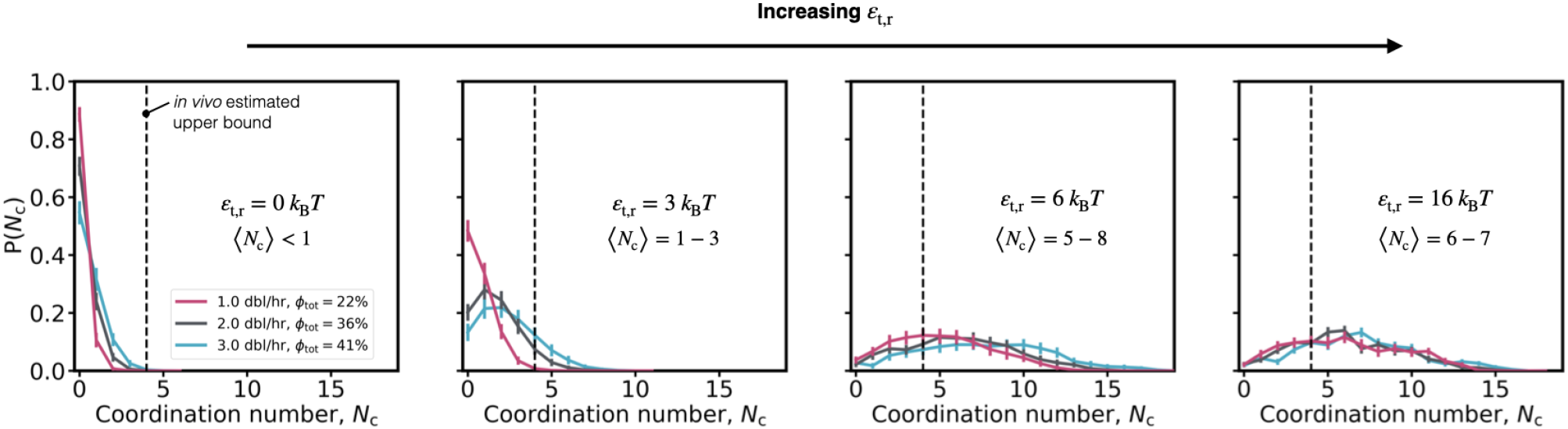
Strong isotropic *ε*_t,r_ *≈* 6 − 16 *k*_B_*T* attractions drive unphysiological coordination numbers of ternary complexes around ribosomes. Coordination number distributions highlight an increase in the number of ternary complexes bound to each ribosome, both with increasing growth rate (red to grey to blue) and attraction strength (left to right). The average number of ternary complex contacts per ribosome, ⟨*N*_c_⟩ , exceeds the *in vivo* estimate of 4 ternary complexes per ribosome (vertical dashed black line, [16]) for attraction strengths *ε*_t,r_ *>* 6 *k*_B_*T* . In our model, isotropic attraction allows a maximum of *N*_c_ = 32 based on close packing of spheres of size *a*_tern_ = 5.9 nm around spheres of size *a*_rib_ = 13 nm [86].

**Figure S4:**
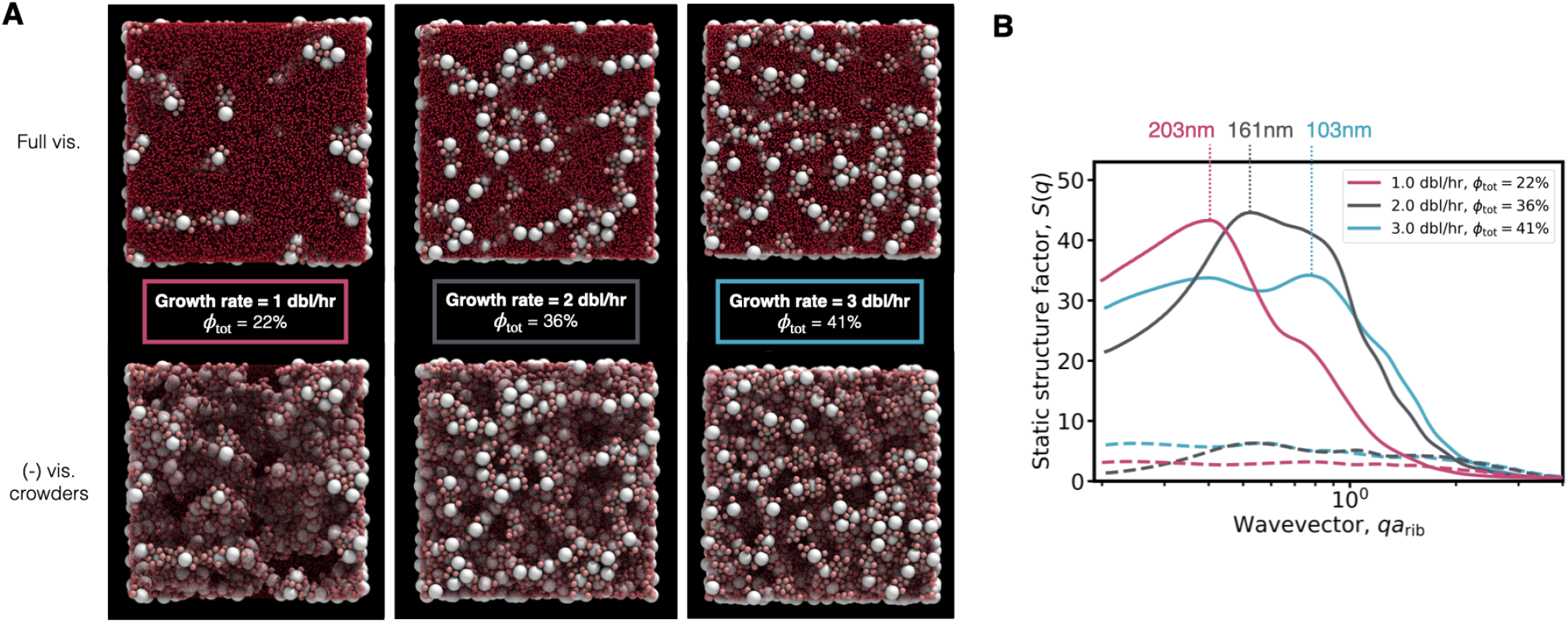
Schematics for modeling physical interactions and chemical reactions in translation voxels of the E. coli cytoplasm in LAMMPS. **(A)** Visualizations of E. coli translation voxels with strong isotropic attractions 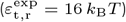 between ternary complexes and ribosomes at three growth rates (left to right). *ϕ* is the total system volume fraction (see ***Methods***), which increases from 22% to 41% as cell growth rate increases. **(Upper)** Full visualization with ribosomes (white), ternary complexes (pink), and crowder proteins (red). **(Lower)** Visualizations with transparent crowder proteins, highlighting the internal voxel microstructure. **(B)** Static structure factor, *S*(*q*), for growth rates of 1 (magenta), 2 (grey), and 3 dbl/hr (cyan) for the hard sphere (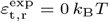, dashed lines) and strongly attractive (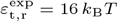, solid lines) cases. The peak, corresponding to the most abundant suspension length scale in real space, is marked with vertical dotted lines. **(C)** Effective cytoplasmic viscosity, *η*_eff,tern_, experienced by ternary complexes in translation voxels (Eq. 10), versus growth rate and volume fraction for the hard sphere (blue) and strongly attractive (red) cases. The viscosity of water, *η*_0_ , at a temperature of 37C (horizontal dashed line) and the estimated *in vivo* viscosity experienced in *E. coli* by a protein with the same molecular weight as a ternary complex (horizontal dotted line [37]) are marked.

**Figure S5:**
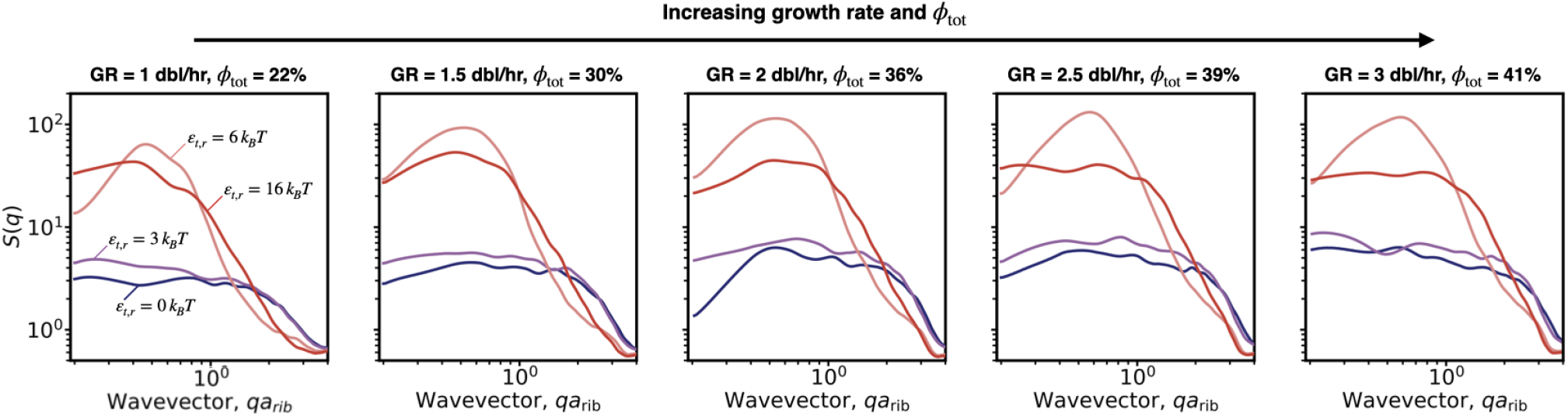
Structural length scales in the *E. coli* cytoplasm. Static structure factor, *S*(*q*), as a function of cellular growth rate and overall volume fraction (left to right, marked at top), and strength of attractions between ternary complexes and ribosomes (colors, marked inset on left). The probability of long-range correlations at small wavevectors is only slightly increased by weak isotropic attractions (*ε*_t,r_ = 3 *k*_B_*T* ), suggesting the formation of transient low-order clusters. A single peak emerges for the *ε*_t,r_ = 6 *k*_B_*T* attractive case, characteristic of a bicontinuous gel created by arrested phase separation. In the *ε*_t,r_ = 16 *k*_B_*T* attractive case, there is an abundance of correlations at longer length-scales (shorter wavevectors), but the stringier structure is not as well characterized by a single length-scale since ongoing phase separation is slowed by strong bonding [67].

**Figure S6:**
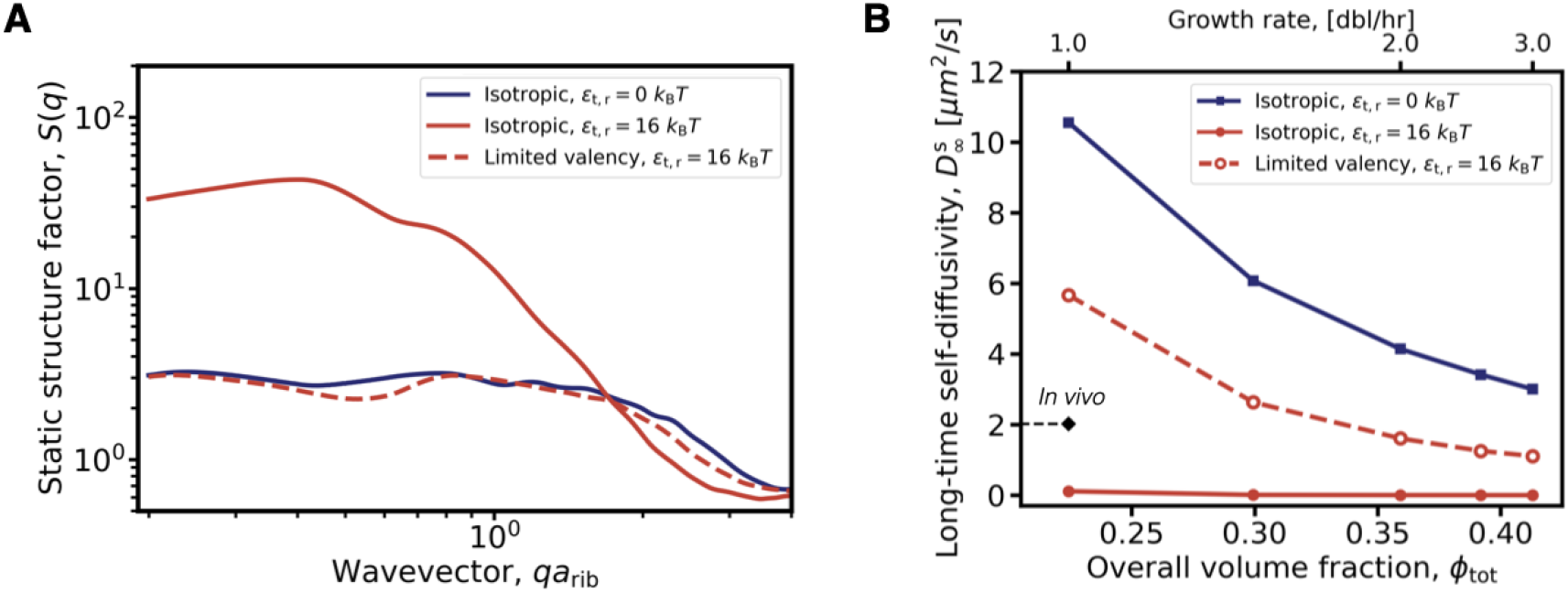
Limiting the valency of attractions between ternary complexes and ribosomes prevents arrested phase separation. **(A)**. Static structure factor, *S*(*q*), at a growth rate of 1 dbl/hr in the purely-repulsive limit (blue, solid line) and with isotropic (red, solid line) and limited valency (red, dashed line) attractions between ternary complexes and ribosomes. Limiting the interaction valency prevents the abundance of longer length-scale correlations observed in the isotropic case. **(B)**. Long-time self-diffusivity of ternary complexes as a function of growth rate and total volume fraction. The experimentally-observed apparent diffusivity is shown for a growth rate of 1.0 dbl/hr (♦) [16]. Limiting the interaction valency improves agreement with the in vivo measurement, without fully arresting dynamics as in the isotropic case. The remaining gap could be attributed to the limited temporal resolution of particle tracking experiments (∼ 2 ms), which are sufficiently large to sample the confinement in a given observation window. We thus might expect experimental measurements (♦) to be artificially low relative to our unbound simulations.

**Figure S7:**
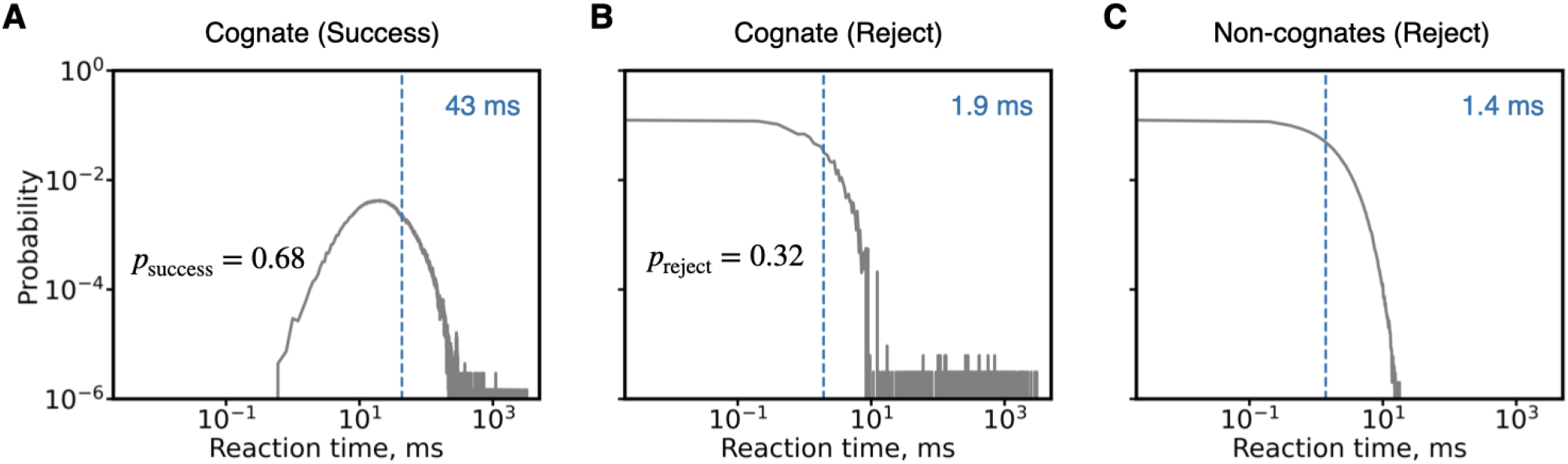
Distribution of time spent by cognate and non-cognate ternary complexes in reactions with ribosomes, computed by sampling kinetic trajectories. **(A)**. Reaction times for cognate ternary complexes that successfully react (with probability *p*_success_ inset, black) and **(B)** those that are rejected (with probability *p*_reject_ inset, black). **(C)** Reaction times for non-cognate ternary complexes, of which all are rejected. In each plot, the average latency is marked (vertical blue dashed line, displayed at top right).

**Figure S8:**
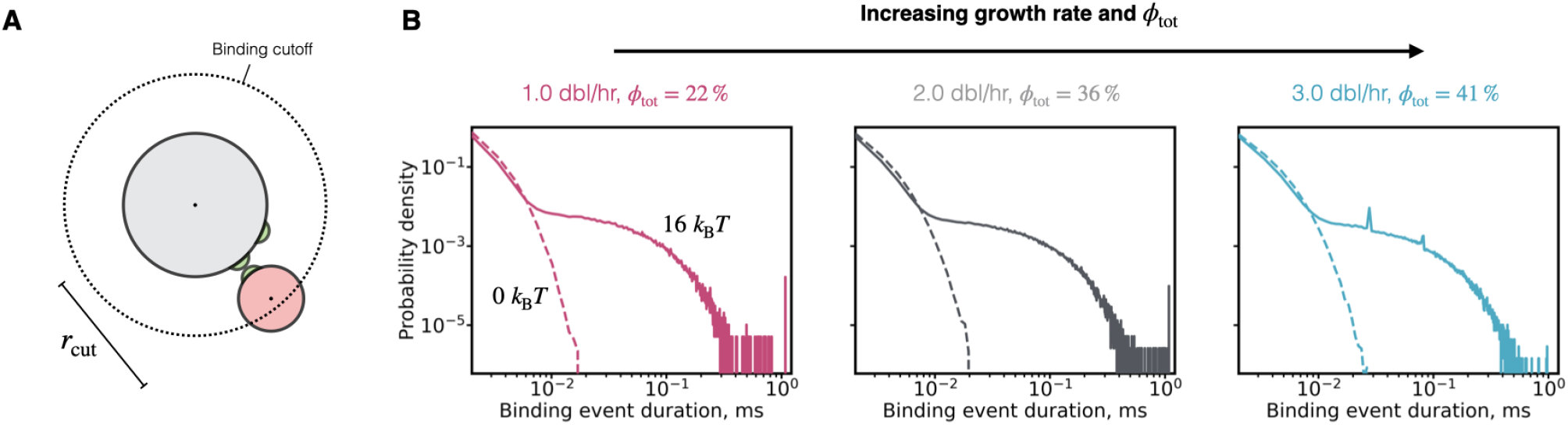
Probability distributions of the duration of physical ternary complex - ribosome binding events. (A). Schematic of binding cutoff between limited valence ternary complexes and ribosomes, chosen to be *r*_cut_ = 2.1 nm based on the radius of the binding site protrusions and width of the attractive potential well. **(B)**. Probability density function of the duration of binding events, defined as when a ternary complex and ribosome come within a bond distance (*r*_cut_). Across all studied growth rates (left to right), purely entropic binding events are brief, lasting at most *∼* 20 µs (dashed lines, 0 *k*_B_*T* ). The strong limited valence case (solid lines, 16 *k*_B_*T* ) show binding times up to 1 ms, on the order of chemical reaction timescales, but below the upper limit of 2 ms inferred from experiment [16].

**Figure S9:**
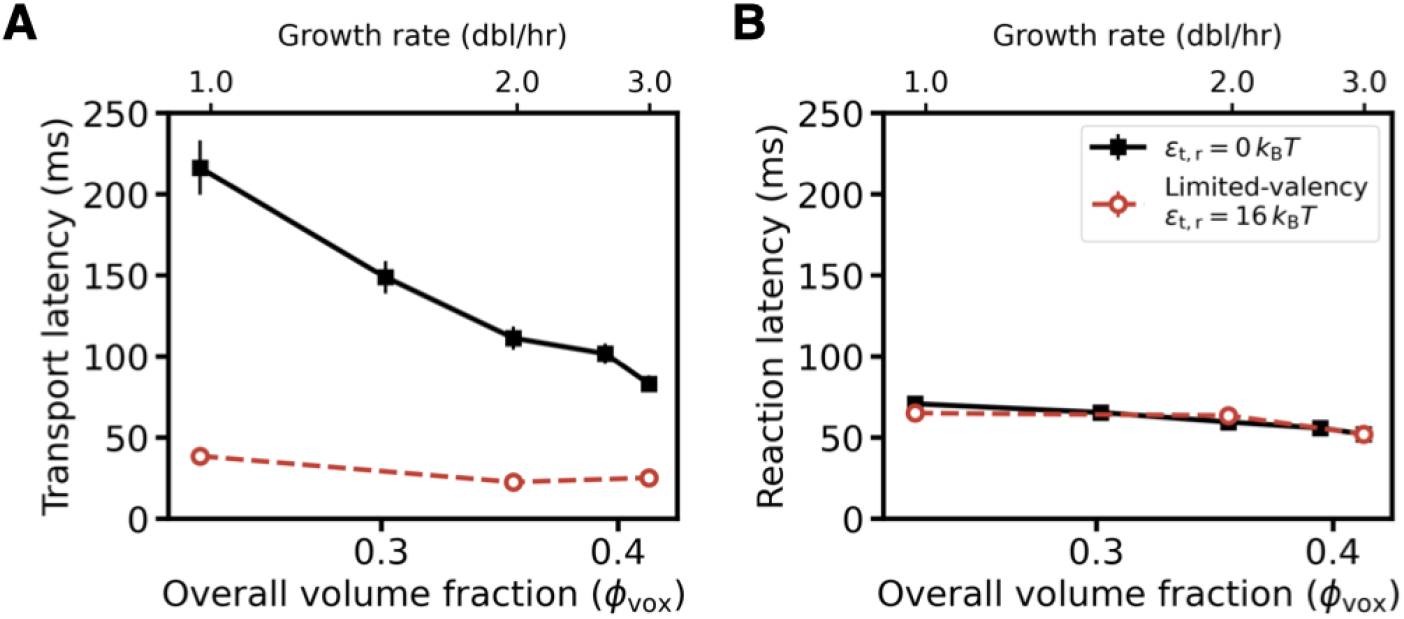
EF-Tu·L12 interactions speed elongation by facilitating faster transport. Simulation results from translation voxels without attractions (black line, [8]) and with limited-valency attractions between ternary complexes and ribosomes (red line, this work). **(A)** Transport latency, which decreases with increasing growth rate (left to right), is reduced by pre-loading of ternary complexes onto ribosomes. **(B)** Reaction latency is unchanged by cognate-independent pre-loading.

**Figure S10:**
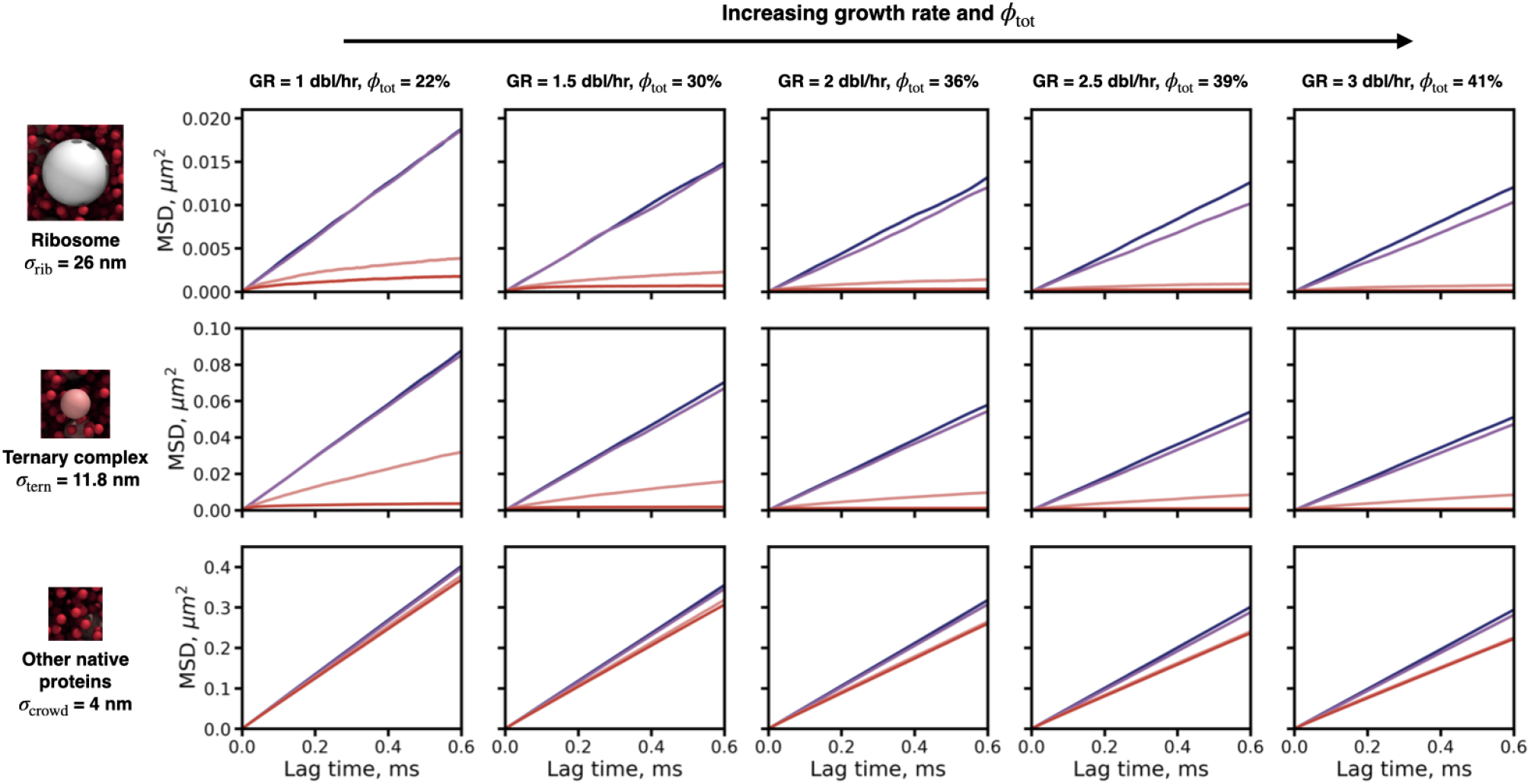
Dynamics of translation molecules are suppressed by increased crowding and attraction strength. Mean squared displacements versus lag time are plotted for ribosomes (top row), ternary complexes (middle row), and other native cytoplasmic proteins (bottom row), for each growth rate (left to right) and attraction strengths of *ε*_t,r_ = 0 *k*_B_*T* (blue), 3 *k*_B_*T* (purple), 6 *k*_B_*T* (pink), and 16 *k*_B_*T* (red).

**Figure S11:**
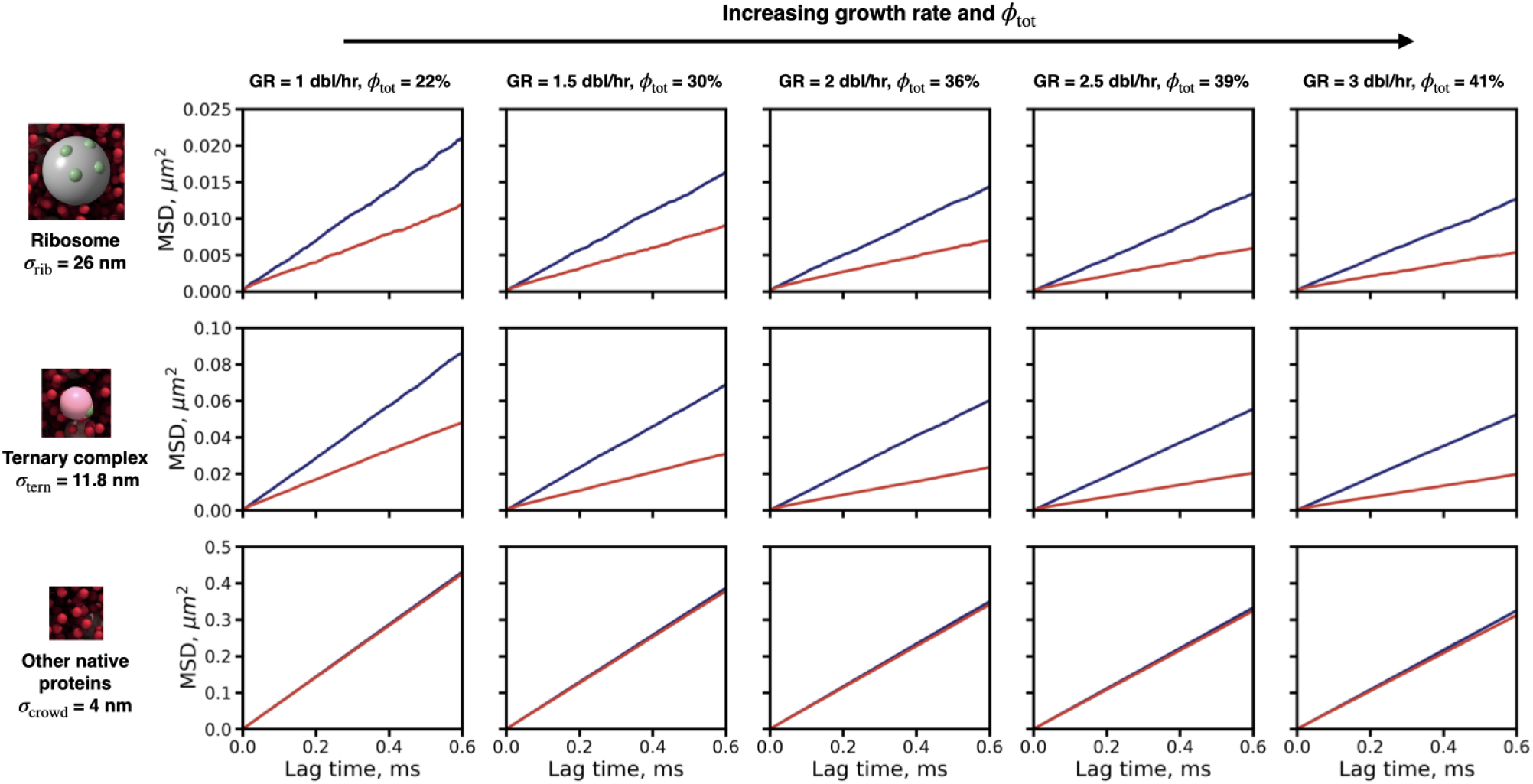
Limiting the valency of attractions between ternary complexes and ribosomes permits more particle mobility across growth rates. Mean squared displacements versus lag time are plotted for ribosomes (top row), ternary complexes (middle row), and other native cytoplasmic proteins (bottom row), for each growth rate (left to right) and limited valency attraction strengths of *ε*_t,r_ = 0 *k*_B_*T* (blue) and *ε*_t,r_ = 16 *k*_B_*T* (red).

**Figure S12:**
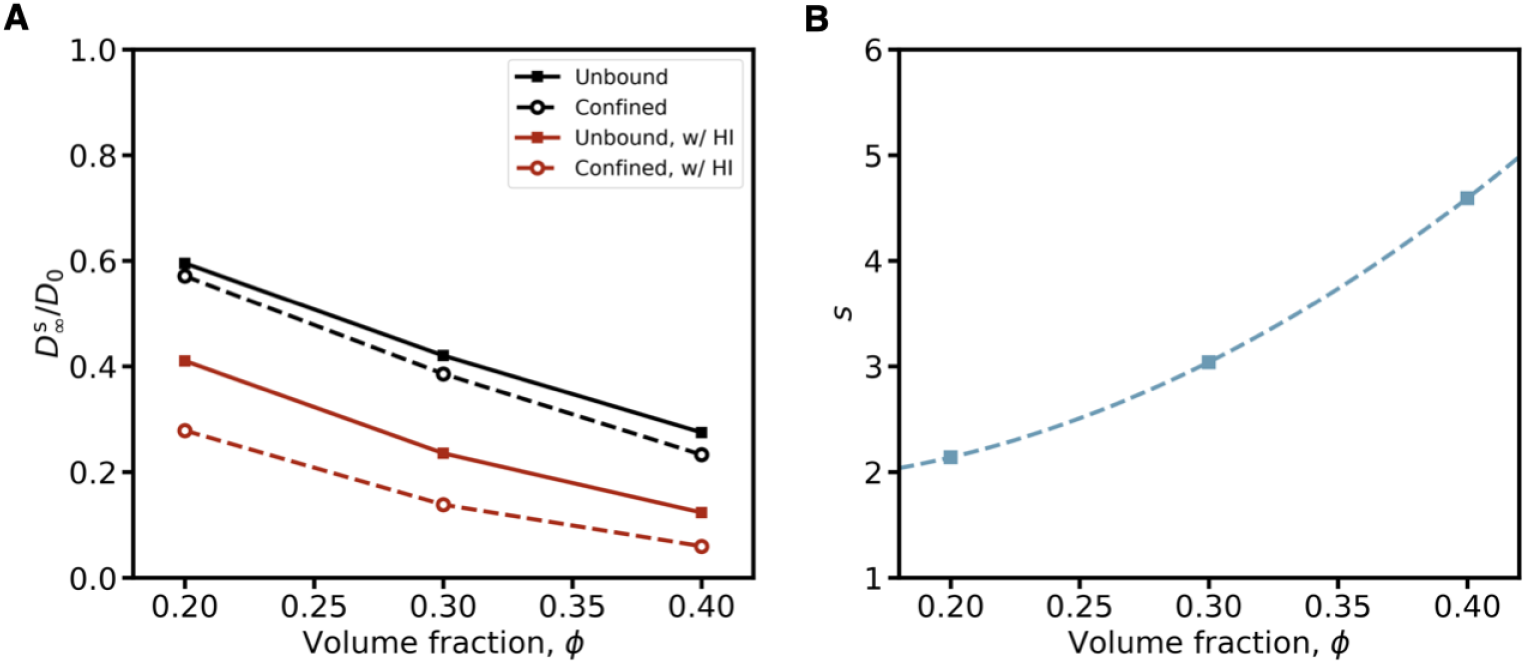
Hydrodynamic interactions and spherical confinement slow diffusion. **(A)** The long-time self-diffusivity, 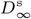, normalized by the lone particle diffusivity, *D*_0_ , in a monodisperse hard-sphere suspension, calculated as a function of volume fraction *ϕ*, the contact value of the radial distribution function g(r), and the short-time self-diffusivity 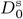, as described in Supplementary Methods . The presence of both confinement (dashed lines) and hydrodynamic interactions (red lines) reduces the expected long-time self-diffusivity. **(B)** The scaling factor, *s*, is calculated via the ratio of 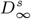 for the unbound and confined + Hydrodynamic Interactions (HI) cases (symbols), and is fit to a second-order polynomial across all volume fractions (dashed line).

**Figure S13:**
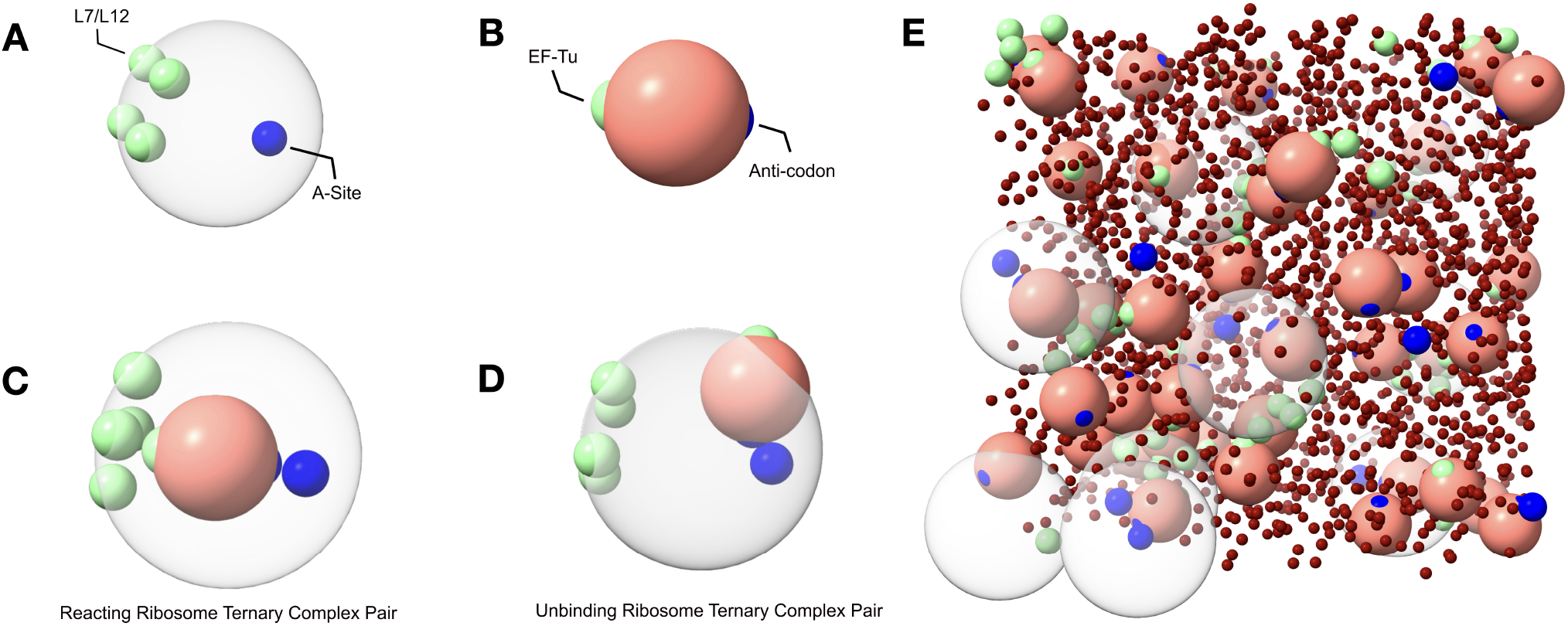
Schematics for modeling physical interactions and chemical reactions in translation voxels of the E. coli cytoplasm in LAMMPS. **(A)** Ribosomes (grey) are represented as rigid bodies with patches representing four L12 subunits (green) and one internal A-site codon (blue). **(B)** Ternary complexes (pink) are represented with as rigid bodies with one EF-Tu (green) and anti-codon patch (blue). **(C)** After a reaction begins, the ternary complex is internalized within the ribosome by turning off the hard sphere repulsion and turning on a strong attraction between the A-site and anticodon. During reactions, other ternary complexes are allowed to occupy the three free L12 subunits but are not allowed to enter the ribosome. **(D)** Following the conclusion of a reaction, the ternary complex is released by a soft Morse repulsion between formerly reacting pairs. **(E)** Thousands of unique translation voxels are simulated to capture the physiological distribution of translation molecules and to calculate reaction, transport, and elongation latencies. Visualizations were created using UCSF ChimeraX [68].

**Figure S14:**
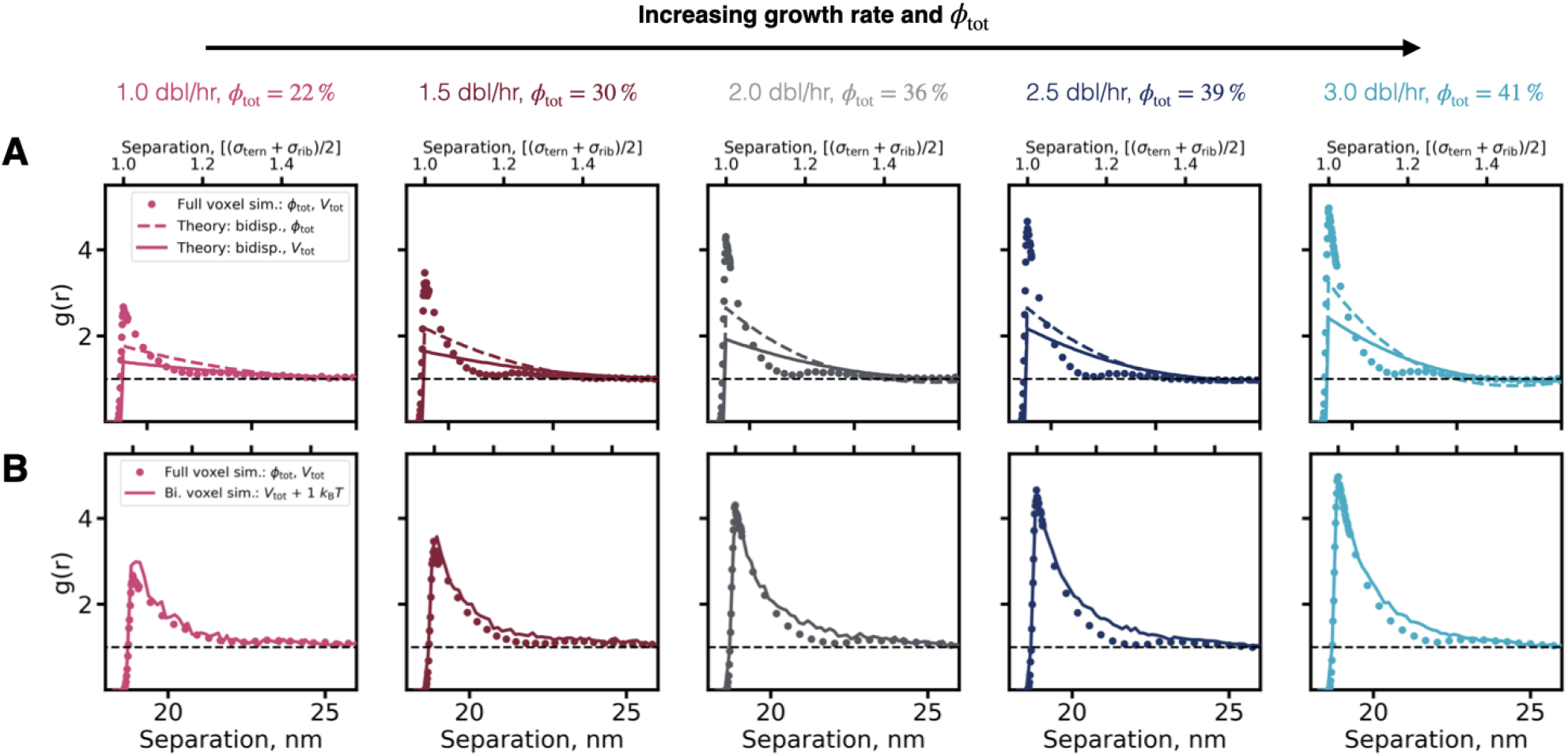
Small proteins in the *E. coli* cytoplasm induce depletion interactions of strength *ε*_t,r_ *≈* 1 *k*_B_*T* between translation molecules. **(A)**. Radial distribution functions between ternary complexes and ribosomes, *g*(*r*), calculated in simulations of translation voxels (symbols), highlight structural deviations from the ideal gas limit (dashed black line) across all growth rates. Analytical radial distribution functions, calculated for bidisperse suspensions of ternary complexes and ribosomes with the same abundances as in the full translation voxels, are also shown for systems with the same *ϕ*_tot_ (colored dashed line) and *V*_tot_ (colored solid line). Volume fraction alone is insufficient to recover the microstructure observed in full cytoplasmic voxels (i.e., the peak at contact, 18.9 nm), suggesting the presence of depletion interactions. **(B)**. Bidisperse systems with added *ε*_t,r_ = 1 *k*_B_*T* Yukawa attractions (solid lines) recover the contact peaks of the tridisperse *g*(*r*), across all growth rates, thus estimating the strength of the depletion interaction.

**Figure S15:**
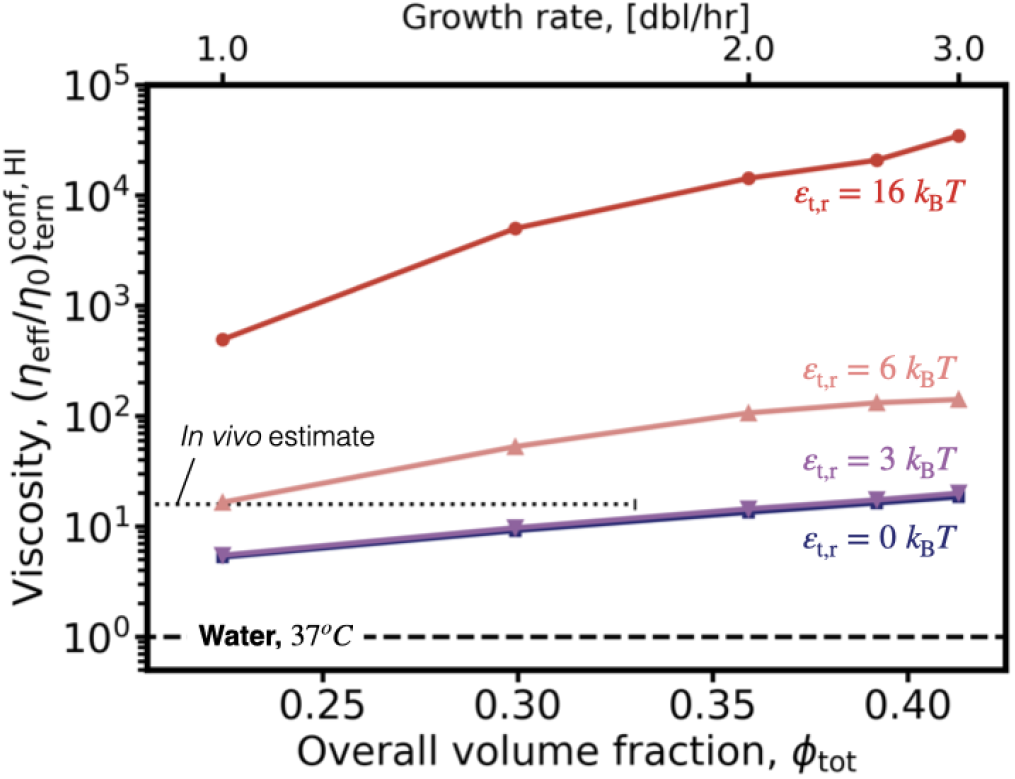
Arrest increases cytoplasmic viscosity beyond *in vivo* estimates. Effective cytoplasmic viscosity experienced by ternary complexes in translation voxels (Eq. 10), versus growth rate and volume fraction for all studied attraction strengths of *ε*_t,r_ = 0 *k*_B_*T* (blue), 3 *k*_B_*T* (purple), 6 *k*_B_*T* (pink), and 16 *k*_B_*T* (red). The viscosity of water, *η*_0_ , at a temperature of 37^*o*^ *C* (horizontal dashed line) and the estimated *in vivo* viscosity experienced by a protein with a molecular weight of 69 kDa (*M*_tern_) in *E. coli* (horizontal dotted line, [37]) are marked.

**Figure S16:**
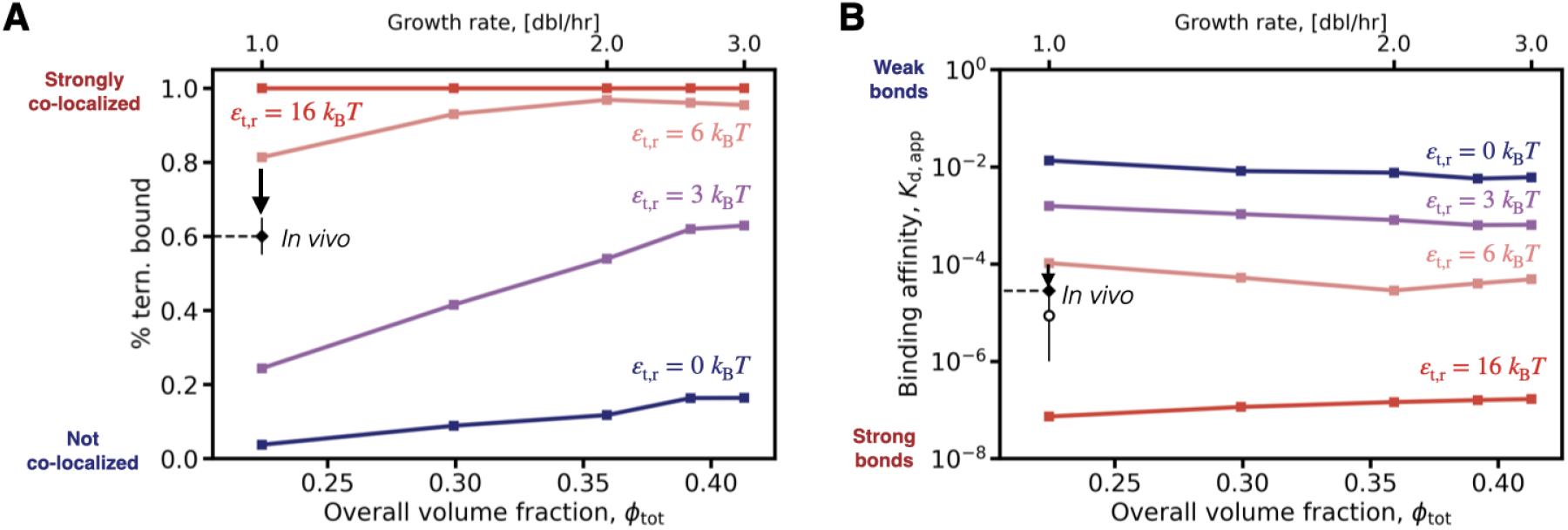
Physical co-localization and biochemical affinity measurements suggest that a limited interaction valency is needed to recover experimental measurements. **(A)** A measure of physical co-localization: the percent of ternary complexes bound to ribosomes for isotropic attraction strengths of *ϵ*_*t*,*r*_ = 0 − 16*k*_b_*T* (colors). Stronger attractions drive more co-localization of translation molecules, and suggest that an isotropic attraction strength less than *ϵ*_*t*,*r*_ = 6*k*_b_*T* would best recover experimental measurements at a growth rate of 1.0 dbl/hr (♦) [16]. **(B)** A measure of biochemical binding affinity: the apparent equilibrium dissociation constant *K*_d,app_ for the interaction between ternary complexes and ribosomes, with isotropic attraction strengths of *ϵ*_*t*,*r*_ = 0 − 16*k*_b_*T* (colors). Stronger attractions drive stronger binding affinities, and suggest that an isotropic attraction strength greater than *ϵ*_*t*,*r*_ = 6*k*_b_*T* would best recover experimental measurements at a growth rate of 1.0 dbl/hr (♦) [16]. Validation of Eq. S4 is marked (*°*), assuming four binding sites on the ribosome, 60% of ternary complexes bound, and abundances and densities of the translation voxel at a growth rate of 1.0 dbl/hr.

**Figure S17:**
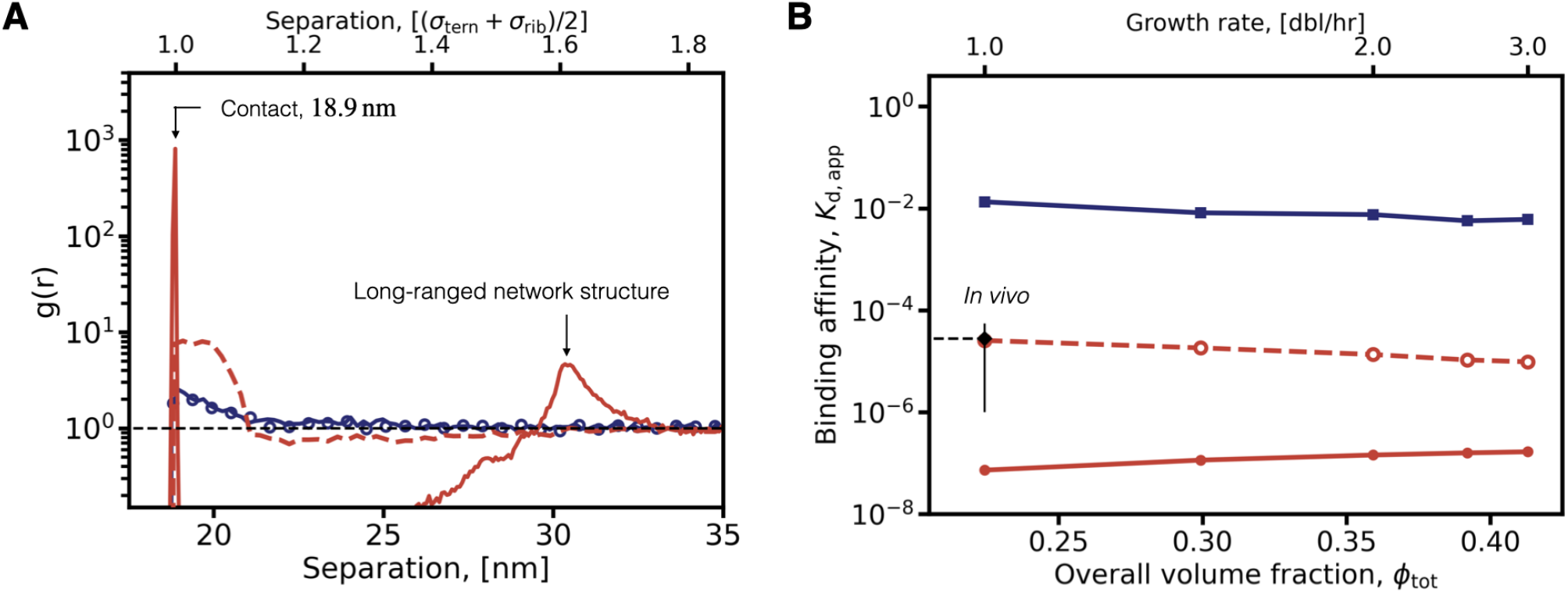
Limited-valency attractions between ternary complexes and ribosomes recover physiological levels of co-localization and binding affinity, while avoiding cytoplasmic arrest. **(A)** Radial distribution functions, *g*(*r*), between ternary complexes and ribosomes at a growth rate of 1 dbl/hr and *ϕ* = 22%. The non-attractive limited-valency case (blue, symbols) quantitatively recovers the microstructure of the non-attractive isotropic case (blue, solid line). Strong isotropic attractions (red solid line) produce a high probability of finding a ternary complex and ribosome at contact, as well as a second peak corresponding with the formation of a long-ranged network structure. Strong limited-valency attractions (red dashed line) increase the contact peak versus the non-attractive case , while avoiding the second peak. **(B)** The apparent equilibrium dissociation constant *K*_d,app_ for the interaction between ternary complexes and ribosomes, as a function of growth rate.

**Figure S18:**
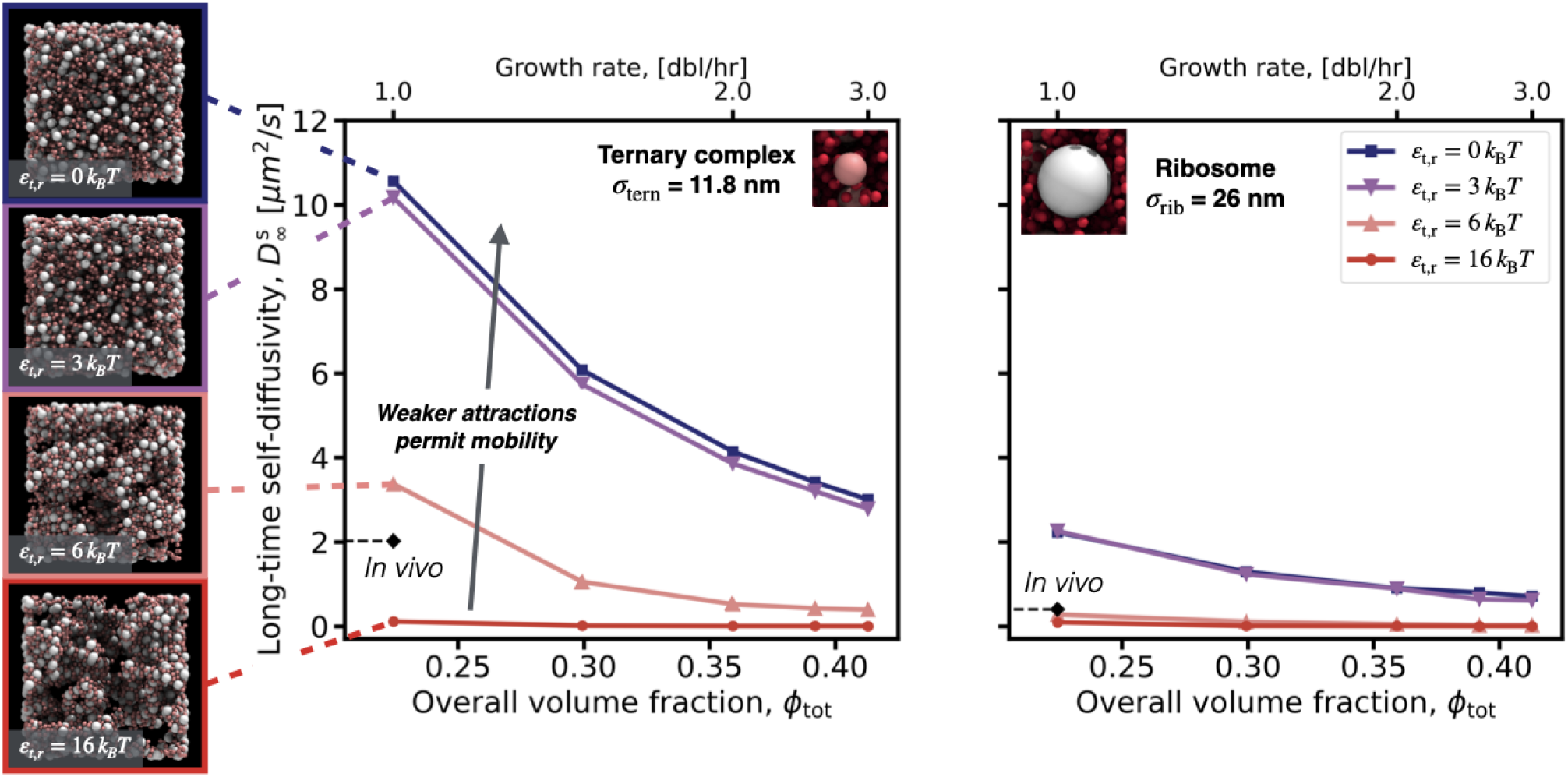
Isotropic attractions weak enough to avoid arrest permit faster diffusion than measured *in vivo*. The long-time self-diffusivity (Eq. 7) for ternary complexes (left) and ribosomes (right) as a function of growth rate and total volume fraction *ϕ*_tot_ , with isotropic attractions of strength *ε*_t,r_ = 0 − 16 *k*_B_*T* (colors, inset legend). The experimentally-observed apparent diffusivity is shown for a growth rate of 1.0 dbl/hr (♦) for both ternary complexes and ribosomes [16]. Simulation screenshots of translation voxels at 1.0 dbl/hr at each studied attraction strength are shown (inset, left).

**Figure S19:**
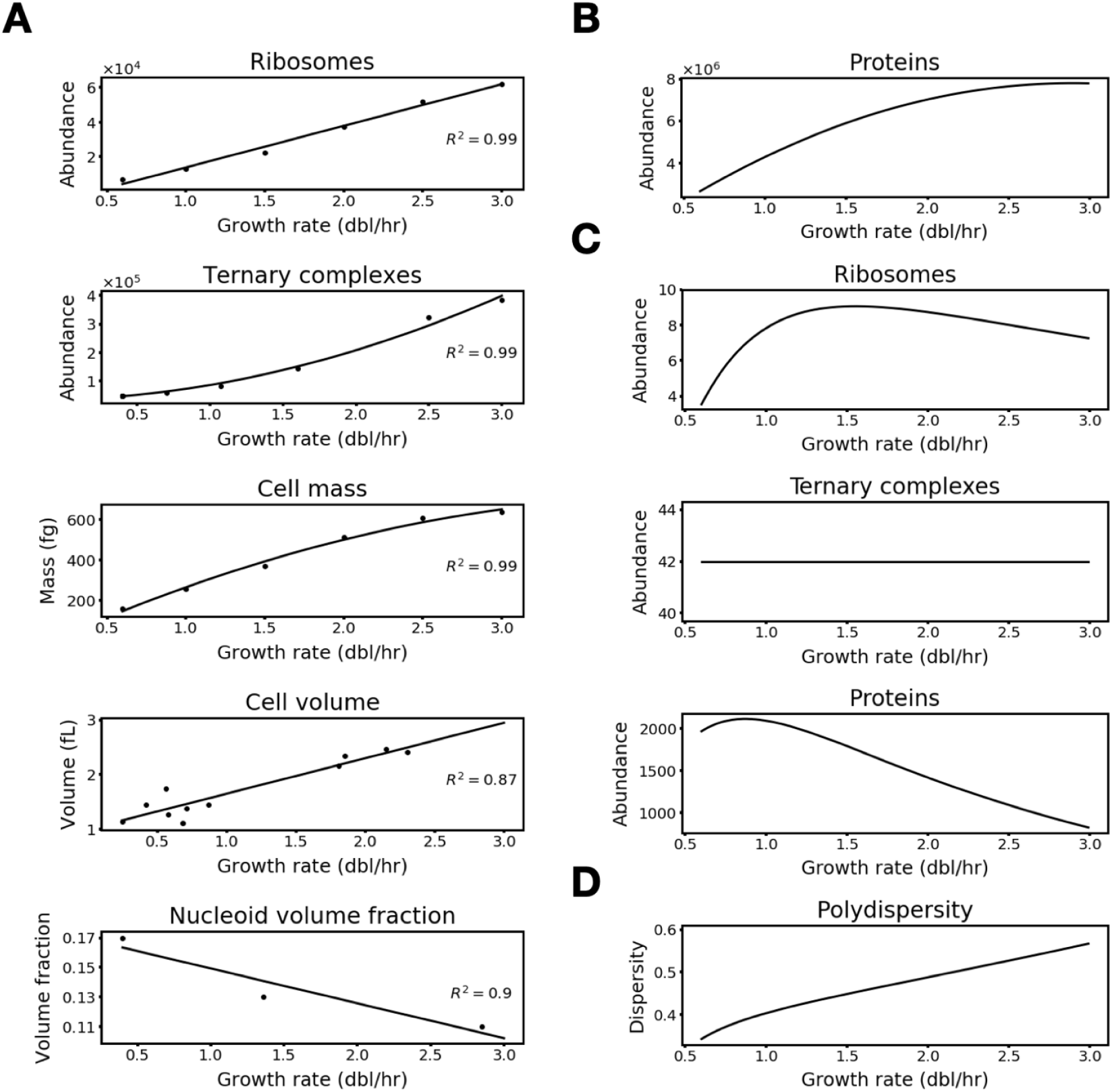
Cell and translation voxel parameters vary with growth rate. **(A)** Polynomial fits of reported cell parameter measurements. **(B)** Estimate of protein abundances in cells across growth rates. **(C)** Estimate of ribosome, ternary complex, and protein abundances in translation voxels across growth rates. **(D)** Estimate of the polydispersity of translation voxels across growth rates. Figure reproduced with permission from [8].

**Figure S20:**
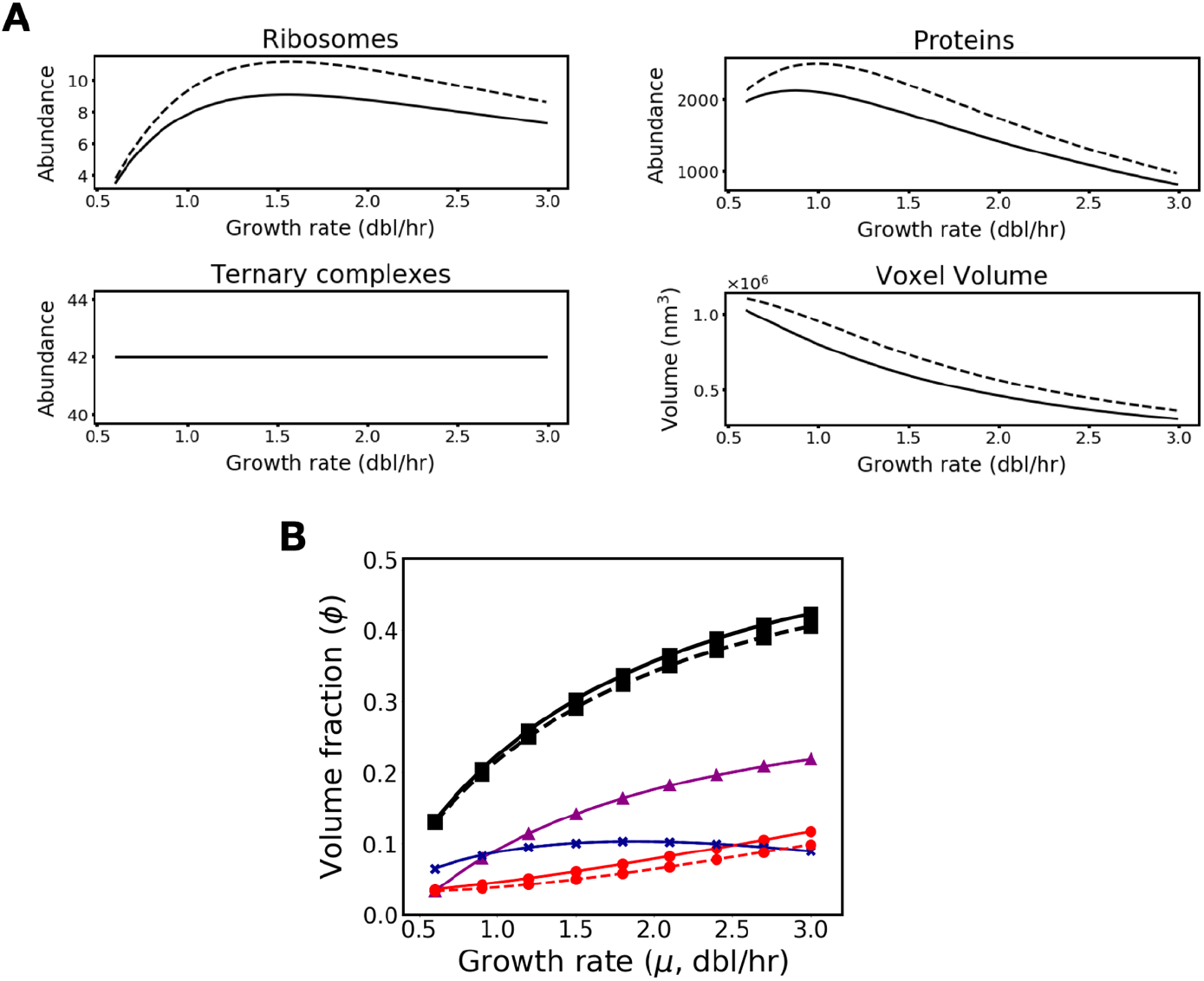
Whether or not literature-determined values for ternary complex abundances account for peptidyl-tRNA does not impact voxel composition or trends. **(A)** Key growth-rate dependent trends in translation voxel composition resulting from the modeling used throughout our manuscript (solid line), in which we interpret ternary complex abundance measurements as not including peptidyl-tRNA, are equivalent to those resulting from assuming that ternary complex abundance measurements do include peptidyl-tRNA (dashed line). **(B)** The volume fraction of ribosomes (purple curve, triangles), ternary complexes (red curve, circles), and proteins (blue curve, x’s), as well as total volume fraction (black curve, squares) are negligibly different for voxels constructed with the assumption that ternary complex measurements do not include peptidyl-tRNA (solid line) and the assumption that they do (dashed line). Figure reproduced with permission from [8].

**Figure S21:**
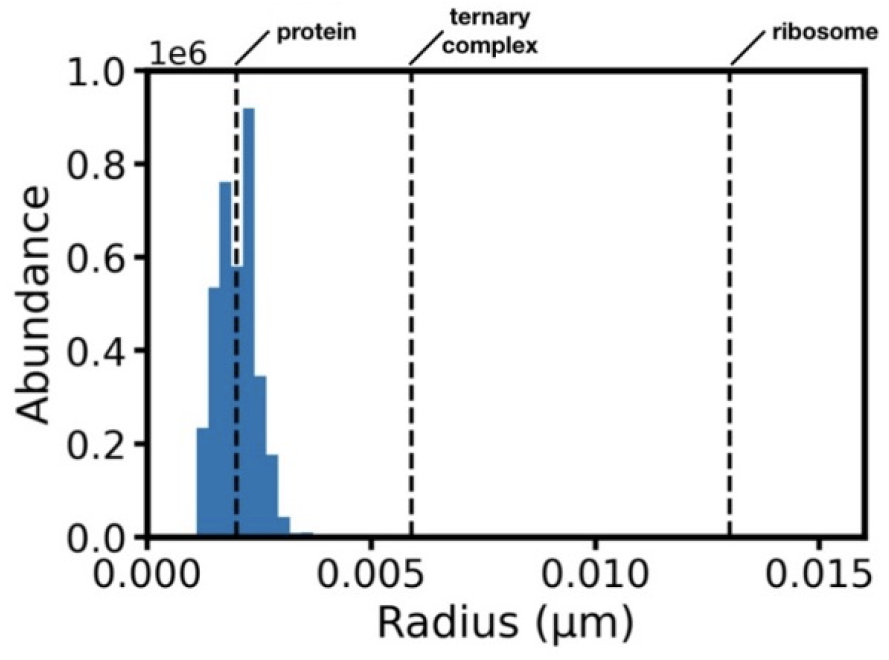
Size distribution of proteins in E. coli. Dashed lines show the size of an average-sized protein, a ternary complex, and a ribosome. Figure reproduced with permission from [8].

## REFERENCES

[1] Ken A. Dill, Kingshuk Ghosh, and Jeremy D. Schmit. Physical limits of cells and proteomes. Proceedings of the National Academy of Sciences of the United States of America, 108:17876–17882, 11 2011.

[2] Nathan M. Belliveau, Griffin Chure, Christina L. Hueschen, Hernan G. Garcia, Jane Kondev, Daniel S. Fisher, Julie A. Theriot, and Rob Phillips. Fundamental limits on the rate of bacterial growth and their influence on proteomic composition. Cell Systems, 12:924–944.e2, 9 2021.

[3] Kirill B. Gromadski and Marina V. Rodnina. Kinetic determinants of high-fidelity trna discrimination on the ribosome. Molecular cell, 13:191–200, 2004.

[4] Kirill B. Gromadski, Tina Daviter, and Marina V. Rodnina. A uniform response to mismatches in codon-anticodon complexes ensures ribosomal fidelity. Molecular cell, 21:369–377, 2006.

[5] Ute Kothe and Marina V. Rodnina. Delayed release of inorganic phosphate from elongation factor tu following gtp hydrolysis on the ribosome. Biochemistry, 45:12767–12774, 2006.

[6] Ingo Wohlgemuth, Corina Pohl, and Marina V. Rodnina. Optimization of speed and accuracy of decoding in translation. The EMBO journal, 29:3701–3709, 2010.

[7] Anneli Borg and Måns Ehrenberg. Determinants of the rate of mrna translocation in bacterial protein synthesis. Journal of molecular biology, 427:1835–1847, 2015.

[8] Akshay J. Maheshwari, Alp M. Sunol, Emma Gonzalez, Drew Endy, and Roseanna N. Zia. Colloidal physics modeling reveals how per-ribosome productivity increases with growth rate in escherichia coli. mBio, 12 2022.

[9] Jonathan R. Karr, Jayodita C. Sanghvi, Derek N. MacKlin, Miriam V. Gutschow, Jared M. Jacobs, Benjamin Bolival, Nacyra Assad-Garcia, John I. Glass, and Markus W. Covert. A whole-cell computational model predicts phenotype from genotype. Cell, 150(2):389–401, 2012.

[10] Jan A Stevens, Fabian Grünewald, PA Marco van Tilburg, Melanie König, Benjamin R Gilbert, Troy A Brier, Zane R Thornburg, Zaida Luthey-Schulten, and Siewert J Marrink. Molecular dynamics simulation of an entire cell. Frontiers in Chemistry, 11:1106495, 2023.

[11] Holger Stark, Marina V. Rodnina, Jutta Rinke-Appel, Richard Brimacombe, Wolfgang Wintermeyer, and Marin Van Heel. Visualization of elongation factor tu on the es-cherichia coli ribosome. Nature 997 89:6649, 389:403–406, 1997.

[12] Ute Kothe, Hans Joachim Wieden, Dagmar Mohr, and Marina V. Rodnina. Interaction of helix d of elongation factor tu with helices 4 and 5 of protein l7/12 on the ribo-some. Journal of Molecular Biology, 2004.

[13] Mihaela Diaconu, Ute Kothe, Frank Schlünzen, Niels Fischer, Jörg M. Harms, Alexander G. Tonevitsky, Holger Stark, Marina V. Rodnina, and Markus C. Wahl. Structural basis for the function of the ribosomal l7/12 stalk in factor binding and gtpase activation. Cell, 2005.

[14] Hirotatsu Imai, Toshio Uchiumi, and Noriyuki Kodera. Direct visualization of translational gtpase factor pool formed around the archaeal ribosomal p-stalk by high-speed afm. Proceedings of the National Academy of Sciences, 117(51):32386–32394, 2020.

[15] Anders Liljas and Suparna Sanyal. The enigmatic ribosomal stalk. Quarterly reviews of biophysics, 51:e12, 2018.

[16] Mainak Mustafi and James C. Weisshaar. Simultaneous binding of multiple ef-tu copies to translating ribosomes in live escherichia coli. mBio, 9, 1 2018.

[17] D.S. Goodsell. Elongation factors. RCSB Protein Data Bank, 9 2006.

[18] D.S. Goodsell. Phytase. RCSB Protein Data Bank, sep 2018.

[19] Hans Bremer and Patrick P. Dennis. Modulation of chemical composition and other parameters of the cell at different exponential growth rates. EcoSal Plus, 3, 2 2008.

[20] Benjamin Volkmer and Matthias Heinemann. Condition-dependent cell volume and concentration of escherichia coli to facilitate data conversion for systems biology modeling. PLOS ONE, 6:e23126, 2011.

[21] Hengjiang Dong, Lars Nilsson, and Charles G. Kurland. Co-variation of trna abundance and codon usage in escherichia coli at different growth rates. Journal of Molecular Biology, 260:649–663, 8 1996.

[22] Steen Pedersen, Phillip L. Bloch, Solvejg Reeh, and Frederick C. Neidhardt. Patterns of protein synthesis in e. coli: a catalog of the amount of 140 individual proteins at different growth rates. Cell, 14:179–190, 5 1978.

[23] Alexander Schmidt, Karl Kochanowski, Silke Vedelaar, Erik Ahrné, Benjamin Volkmer, Luciano Callipo, Kèvin Knoops, Manuel Bauer, Ruedi Aebersold, and Matthias Heinemann. The quantitative and condition-dependent escherichia coli proteome. Nature Biotechnology 2 5 4: , 34:104–110, 1 2016.

[24] Conrad L Woldringh and Nanne Nanninga. Structure of nucleoid and cytoplasm of the intact cell, 1985.

[25] Benjamin Volkmer and Matthias Heinemann. Condition-dependent cell volume and concentration of escherichia coli to facilitate data conversion for systems biology modeling. PLoS One, 6:e15727, 2011.

[26] Alexander Schmidt, Karl Kochanowski, Silke Vedelaar, Erik Ahrné, Benjamin Volkmer, Luciano Callipo, Kévin Knoops, Mikael Bauer, Ruedi Aebersold, and Matthias Heinemann. The quantitative and condition-dependent escherichia coli proteome. Nature Biotechnology, 34:104–110, 2015.

[27] Jessica L. Radzikowski, Silke Vedelaar, David Siegel, Álvaro Dario Ortega, Alexander Schmidt, and Matthias Heinemann. Bacterial persistence is an active σs stress response to metabolic flux limitation. Molecular Systems Biology, 12:882, 2016.

[28] Conrad L. Woldringh and Nanne Nanninga. Structure of nucleoid and cytoplasm of the intact cell. In Molecular cytology of Escherichia coli, pages 161–197. Academic Press, 1985.

[29] Hans Bremer and Patrick P. Dennis. Modulation of chemical composition and other parameters of the cell at different exponential growth rates. EcoSal Plus, 3, 2 2008.

[30] Hongping Dong, Lennart Nilsson, and Charles G. Kurland. Co-variation of trna abundance and codon usage in escherichia coli at different growth rates. Journal of Molecular Biology, 260:649–663, 1996.

[31] Barbara S. Schuwirth, Maria A. Borovinskaya, Cathy W. Hau, Wen Zhang, Antón Vila-Sanjurjo, James M. Holton, and Jamie H.Doudna Cate. Structures of the bacterial ribosome at 3.5 Å resolution. Science, 310:827–834, 2005.

[32] Poul Nissen, Søren Thirup, Morten Kjeldgaard, and Jens Nyborg. The crystal structure of Cys-tRNA(Cys)-EF-Tu-GDPNP reveals general and specific features in the ternary complex and in tRNA. Structure, 7(2):143–156, 1999.

[33] Sophia Rudorf, Manuel Thommen, Marina V. Rodnina, and Reinhard Lipowsky. Deducing the kinetics of protein synthesis in vivo from the transition rates measured in vitro. PLoS computational biology, 10:e1003475, 2014.

[34] S Pedersen. Patterns of protein synthesis in e. coli: a catalog of the amount of 140 individual proteins at different growth rates. Cell, 14:179–190, 5 1978.

[35] Ruth A Vanbogelen, Pushpam Sankar, Robert L Clark, Jacqueline A Bogan, and Frederick C Neidhardt. The gene-protein database of escherichia coli: edition 5. Electrophoresis, 13(1):1014–1054, 1992.

[36] Hans Joachim Wieden, Wolfgang Wintermeyer, and Marina V. Rodnina. A common structural motif in elongation factor ts and ribosomal protein l7/12 may be involved in the interaction with elongation factor tu. Journal of Molecular Evolution 2 52:2, 52:129–136, 2001.

[37] Wojciech M. Smigiel, Luca Mantovanelli, Dmitrii S. Linnik, Michiel Punter, Jakob Silberberg, Limin Xiang, Ke Xu, and Bert Poolman. Protein diffusion in escherichia coli cytoplasm scales with the mass ofthe complexes and is location dependent. Science Advances, 8:5387, 8 2022.

[38] G. K. Batchelor. Brownian diffusion of particles with hydrodynamic interaction. J. Fluid Mech., 74(1):1–29, mar 1976.

[39] Christian Aponte-Rivera, Yu Su, and Roseanna N. Zia. Equilibrium structure and diffusion in concentrated hydrodynamically interacting suspensions confined by a spherical cavity. Journal of Fluid Mechanics, 836:413–450, 2 2018.

[40] Emma Gonzalez, Christian Aponte-Rivera, and Roseanna N. Zia. Impact of polydispersity and confinement on diffusion in hydrodynamically interacting colloidal suspensions. J. Fluid Mech., 925:35, 2021.

[41] Alp M. Sunol and Roseanna N. Zia. Confined Brownian suspensions: Equilibrium diffusion, thermodynamics, and rheology. J. Rheol. (N. Y. N. Y)., 67(2):433, jan 2023.

[42] Jes Forchhammer and Lasse Lindahl. Growth rate of polypeptide chains as a function of the cell growth rate in a mutant of escherichia coli 15. Journal of Molecular Biology, 55:563–568, 2 1971.

[43] D. G. Dalbow and R. Young. Synthesis time of β-galactosidase in escherichia coli b/r as a function of growth rate. Biochemical Journal, 150:13–20, 7 1975.

[44] R. Young and H. Bremer. Polypeptide-chain-elongation rate in escherichia coli b/r as a function of growth rate. Biochemical Journal, 160:185–194, 11 1976.

[45] S. Pedersen. Escherichia coli ribosomes translate in vivo with variable rate. The EMBO Journal, 3:2895–2898, 12 1984.

[46] Daphna Frenkiel-Krispin, Smadar Levin-Zaidman, Eyal Shimoni, Sharon G. Wolf, Ellen J. Wachtel, Talmon Arad, Steven E. Finkel, Roberto Kolter, and Abraham Minsky. Regulated phase transitions of bacterial chromatin: A non-enzymatic pathway for generic dna protection. EMBO Journal, 20:1184–1191, 3 2001.

[47] Hans Bremer and Patrick P Dennis. Modulation of chemical composition and other parameters of the cell by growth rate, 1996.

[48] Stefan Klumpp, Matthew Scott, Steen Pedersen, and Terence Hwa. Molecular crowding limits translation and cell growth. Proceedings of the National Academy of Sciences of the United States of America, 110:16754–16759, 10 2013.

[49] Jake L. Weissman, Shengwei Hou, and Jed A. Fuhrman. Estimating maximal microbial growth rates from cultures, metagenomes, and single cells via codon usage patterns. Proc. Natl. Acad. Sci. U. S. A., 118(12), mar 2021.

[50] Iakov I. Davydov, Ingo Wohlgemuth, Irena I. Artamonova, Henning Urlaub, Alexander G. Tonevitsky, and Marina V. Rodnina. Evolution of the protein stoichiometry in the L12 stalk of bacterial and organellar ribosomes. Nat. Commun., 4, 2013.

[51] Chandra Sekhar Mandava, Kristin Peisker, Josefine Ederth, Ranjeet Kumar, Xueliang Ge, Witold Szaflarski, and Suparna Sanyal. Bacterial ribosome requires multiple L12 dimers for efficient initiation and elongation of protein synthesis involving IF2 and EF-G. Nucleic Acids Res., 40(5):2054–2064, mar 2012.

[52] Rebecca M. Voorhees and V. Ramakrishnan. Structural basis of the translational elongation cycle. 10.1146/annurev-biochemx9-92, 82:203– 236, 6 2013.

[53] Irmantas Mogila, Giedre Tamulaitiene, Konstanty Keda, Albertas Timinskas, Audrone Ruksenaite, Giedrius Sasnauskas, Česlovas Venclovas, Virginijus Siksnys, and Gintautas Tamulaitis. Ribosomal stalk-captured carf-rele ribonuclease inhibits translation following crispr signaling. Science, 382(6674):1036–1041, 2023.

[54] H. J. Grosjean, S. De Henau, and D. M. Crothers. On the physical basis for ambiguity in genetic coding interactions. Proc. Natl. Acad. Sci., 75(2):610–614, feb 1978.

[55] Declan A Gray, Gaurav Dugar, Pamela Gamba, Henrik Strahl, Martijs J Jonker, and Leendert W Hamoen. Extreme slow growth as alternative strategy to survive deep starvation in bacteria. Nature communications, 10(1):890, 2019.

[56] Zane R. Thornburg, David M. Bianchi, Troy A. Brier, Benjamin R. Gilbert, Tyler M. Earnest, Marcelo C.R. Melo, Nataliya Safronova, James P. Sáenz, András T. Cook, Kim S. Wise, Clyde A. Hutchison, Hamilton O. Smith, John I. Glass, and Zaida Luthey-Schulten. Fundamental behaviors emerge from simulations of a living minimal cell. Cell, 185:345–360.e28, 1 2022.

[57] Derek N. Macklin, Travis A. Ahn-Horst, Heejo Choi, Nicholas A. Ruggero, Javier Carrera, John C. Mason, Gwanggyu Sun, Eran Agmon, Mialy M. DeFelice, Inbal Maayan, Keara Lane, Ryan K. Spangler, Taryn E. Gillies, Morgan L. Paull, Sajia Akhter, Samuel R. Bray, Daniel S. Weaver, Ingrid M. Keseler, Peter D. Karp, Jerry H. Morrison, and Markus W. Covert. Simultaneous cross-evaluation of heterogeneous {E. coli} datasets via mechanistic simulation. Science, 369:eaav3751, 7 2020.

[58] Aidan P. Thompson, H. Metin Aktulga, Richard Berger, Dan S. Bolintineanu, W. Michael Brown, Paul S. Crozier, Pieter J. in ‘t Veld, Axel Kohlmeyer, Stan G. Moore, Trung Dac Nguyen, Ray Shan, Mark J. Stevens, Julien Tranchida, Christian Trott, and Steven J. Plimpton. Lammps - a flexible simulation tool for particle-based materials modeling at the atomic, meso, and continuum scales. Computer Physics Communications, 271:108171, 2 2022.

[59] Roseanna N Zia, Benjamin J Landrum, and William B Russel. A micro-mechanical study of coarsening and rheology of colloidal gels: Cage building, cage hopping, and smoluchowski’s ratchet. Journal of Rheology, 58:1121, 2014.

[60] Jan Spitzer and Bert Poolman. The role of biomacromolecular crowding, ionic strength, and physicochemical gradients in the complexities of life’s emergence. Microbiology and Molecular Biology Reviews, 73:371–388, 2009.

[61] Rico F Tabor, Chu Wu, Franz Grieser, Raymond R Dagastine, and Derek Y C Chan. Measurement of the hydrophobic force in a soft matter system. J. Phys. Chem. Lett, 4:3877, 2013.

[62] Benjamin Folch, Marianne Rooman, and Yves Dehouck. Thermostability of salt bridges versus hydrophobic interactions in proteins probed by statistical potentials. J. Chem. Inf. Model., 48:119–127, 2008.

[63] D. Eric Anderson, Wayne J. Becktel, and F. W. Dahlquist. ph-induced denaturation of proteins: A single salt bridge contributes 3-5 kcal/mol to the free energy of folding of t4 lysozyme. Biochemistry, 29:2403–2408, 3 1990.

[64] Liguo Jiang, Siqin Cao, Peter Pak Hang Cheung, Xiaoyan Zheng, Chris Wai Tung Leung, Qian Peng, Zhigang Shuai, Ben Zhong Tang, Shuhuai Yao, and Xuhui Huang. Real-time monitoring of hydrophobic aggregation reveals a critical role of cooperativity in hydrophobic effect. Nature Communications 2 7 8: , 8:1–8, 5 2017.

[65] Marie Leijonmarck and Anders Liljas. Structure of the c-terminal domain of the ribosomal protein l7l12 from escherichia coli at 1.7 å. Journal of Molecular Biology, 195:555–579, 6 1987.

[66] Jennifer L. Hofmann, Akshay J. Maheshwari, Alp M. Sunol, Drew Endy, and Roseanna N. Zia. Ultra-weak protein-protein interactions can modulate proteome-wide searching and binding. bioRxiv, page 2022.09.30.510365, 10 2022.

[67] Lilian C. Johnson, Roseanna N. Zia, Esmaeel Moghimi, and George Petekidis. Influence of structure on the linear response rheology of colloidal gels. Journal of Rheology, 63:583, 5 2019.

[68] Eric F. Pettersen, Thomas D. Goddard, Conrad C. Huang, Elaine C. Meng, Gregory S. Couch, Tristan I. Croll, John H. Morris, and Thomas E. Ferrin. Ucsf chimerax: Structure visualization for researchers, educators, and developers. Protein Science, 30:70–82, 1 2021.

[69] Paul Langevin. Sur la theorie du mouvement brownien. Comptes-rendus de l’Académie des sciences, 146:530–533, 1908.

[70] L. Martinez, R. Andrade, E. G. Birgin, and J. M. Martínez. PACKMOL: A package for building initial configurations for molecular dynamics simulations. J. Comput. Chem., 30(13):2157–2164, oct 2009.

[71] Michael P. Allen and Dominic J. Tildesley. Computer Simulation of Liquids. Oxford University Press, 2nd editio edition, 11 2017.

[72] Axel Brünger, Charles L. Brooks, and Martin Karplus. Stochastic boundary conditions for molecular dynamics simulations of st2 water. Chemical Physics Letters, 105:495– 500, 3 1984.

[73] William Humphrey, Andrew Dalke, and Klaus Schulten. Vmd: Visual molecular dynamics. Journal of Molecular Graphics, 14:33–38, 2 1996.

[74] C.W.J Beenakker and P. Mazur. Diffusion of spheres in a concentrated suspension ii. Physica, 126A, 1984.

[75] John F. Brady. The long-time self-diffusivity in concentrated colloidal dispersions. Journal of Fluid Mechanics, 272:109–134, 1994.

[76] George Gabriel Stokes. On the effect of the internal friction of fluids on the motion of pendulums. Transactions of the Cambridge Philosophical Society, 9, 1851.

[77] A. Einstein. Über die von der molekularkinetischen theorie der wärme geforderte bewegung von in ruhenden flüssigkeiten suspendierten teilchen. Annalen der Physik, 322:549–560, 1 1905.

[78] Roseanna N Zia. Active and Passive Microrheology: Theory and Simulation. Annu. Rev. Fluid Mech, 50:371–405, 2018.

[79] Donald A McQuarrie. Statistical mechanics.Harper & Row, New York, 1976.

[80] Joshua S. Madin, Daniel A. Nielsen, Maria Brbic, Ross Corkrey, David Danko, Kyle Edwards, Martin K.M. Engqvist, Noah Fierer, Jemma L. Geoghegan, Michael Gillings, Nikos C. Kyrpides, Elena Litchman, Christopher E. Mason, Lisa Moore, Søren L. Nielsen, Ian T. Paulsen, Nathan D. Price, T. B.K. Reddy, Matthew A. Richards, Eduardo P.C. Rocha, Thomas M. Schmidt, Heba Shaaban, Maulik Shukla, Fran Supek, Sasha G. Tetu, Sara Vieira-Silva, Alice R. Wattam, David A. Westfall, and Mark Westoby. A synthesis of bacterial and archaeal phenotypic trait data. Sci. Data 2 2 7 , 7(1):1–8, jun 2020.

[81] Jerome K. Percus and George J. Yevick. Analysis of Classical Statistical Mechanics by Means of Collective Coordinates. Phys. Rev., 110(1):1, apr 1958.

[82] W R Smith, D J Henderson, P J Leonard, J A Barker, and E W Grundke. Fortran codes for the correlation functions of hard sphere fluids. Mol. Phys., 106(1):3–7, 2008.

[83] Loup Verlet and Jean Jacques Weis. Equilibrium Theory of Simple Liquids. Phys. Rev. A, 5(2):939, feb 1972.

[84] E. W. Grundke and D. Henderson. Distribution functions of multi-component fluid mixtures of hard spheres. Mol. Phys., 24(2):269–281, aug 2006.

[85] Hans Bisswanger. Multiple Equilibria, Principles, and Derivations. In Enzym. Kinet., pages 1–26. John Wiley & Sons, Ltd, jun 2017.

[86] B W Clare and D L Kepert. The closest packing of equal circles on a sphere. Proc. R. Soc. London. A. Math. Phys. Sci., 405(1829):329–344, jun 1986.

[87] Jean Pierre Hansen and Ian R. McDonald. Theory of simple liquids. Elsevier Academic Press, 3rd ed. edition, 2006.

[88] Alice L. Thorneywork, Roland Roth, Dirk G.A.L. Aarts, and Roel P.A. Dullens. Communication: Radial distribution functions in a two-dimensional binary colloidal hard sphere system. The Journal of Chemical Physics, 140:161106, 4 2014.

